# From individual behaviors to collective outcomes: fruiting body formation in *Dictyostelium* as a group-level phenotype

**DOI:** 10.1101/2022.04.12.487948

**Authors:** Jennie F. Kuzdzal-Fick, Armando Moreno, Cathleen M.E. Broersma, Tim F. Cooper, Elizabeth A. Ostrowski

**Affiliations:** Department of Biology and Biochemistry, University of Houston, Houston, Texas; School of Natural Sciences, Massey University, Auckland, New Zealand

**Keywords:** collective behaviors, genotype-by-genotype interactions (*G*×*G*), epistasis, cooperation, conflict, joint phenotype

## Abstract

Collective phenotypes, which arise from the interactions among individuals, can be important for the evolution of higher levels of biological organization. However, how a group’s composition determines its collective phenotype remains poorly understood. When starved, cells of the social amoeba *Dictyostelium discoideum* cooperate to build a multicellular fruiting body, and the morphology of the fruiting body is likely advantageous to the surviving spores. We assessed how the number of strains, as well as their genetic and geographic relationships to one another, impact the group’s morphology and productivity. We find that some strains consistently enhance or detract from the productivity of their groups, regardless of the identity of the other group members. We also detect extensive pairwise and higher-order genotype interactions, which collectively have a large influence on the group phenotype. Whereas previous work in *Dictyostelium* has focused almost exclusively on whether spore production is equitable when strains cooperate to form multicellular fruiting bodies, our results suggest a previously unrecognized impact of chimeric co-development on the group phenotype. Our results demonstrate how interactions among members of a group influence collective phenotypes and how group phenotypes might in turn impact selection on the individual.

## Introduction

Individuals frequently form groups and engage in cooperative behaviors within them that benefit the members. In some cases, group benefits arise from a higher-level phenotype that emerges from the interactions among its members. For example, herding behaviors can produce an optical illusion that confuses predators and thereby protects the individuals (Olson et al. 2013). Similarly, multicellularity can be advantageous if large cell clusters render the collective resistant to predation (discussed in Bourke 2011; Herron et al. 2019), external stressors (Kuzdzal-Fick et al. 2019), or allow access to a new niche (Rainey and Travisano 1998). Despite many benefits to group formation, the composition of a group—in terms of the number and identity of cooperating individuals—can impact whether these benefits are realized, resulting in a feedback between group composition and individual fitness (Farine et al. 2015).

Prior studies have shown diverse effects of the number and identity of individuals in a group on collective phenotypes. On one hand, genetic diversity sometimes confers benefits, referred to as *social heterosis* (Nonacs and Kapheim 2008). For example, in ants, increased genetic diversity can increase colony growth rate (Cole and Wiernasz 1999; Modlmeier et al. 2012), improve foraging and nest maintenance (Saar et al. 2018), and increase resistance to disease (Hughes and Boomsma 2004). However, genetically diverse groups can also experience enhanced conflict owing to low relatedness, which can enhance within-group competition and favor selfish behaviors that are detrimental to the group (Keller and Chapuisat 1999; Bourke 2011). Thus, both theory and evidence suggest that the genetic relationships among individuals in a group can impact its collective performance, but the direction of these changes is not easily predicted.

Geographic relationships between component individuals might also influence group performance. For example, selection may favor compatible groupings, leading to patterns of local adaptation, where individuals from the same population are highly productive with one another, but groups composed of individuals from different populations show poor group performance. In animals, conflict can sometimes depend on familiarity, which will often be related to geography. For example, the ‘dear enemy effect’ (DEE) refers to a behavioral phenomenon whereby familiar individuals show reduced aggression, likely because territoriality can confer a fitness cost in the form of injuries, potentially to both parties (Temeles 1994; Christensen and Radford 2018). The opposite phenomenon, the ‘nasty neighbor effect’ (NNE), occurs when neighboring individuals exhibit elevated aggression, most likely because they recognize and compete most strongly with one another. Thus, neighbors can show either enhanced or reduced conflict, and the existence of either the DEE or NNE can provide important insights into the scale of competition.

The social amoeba *Dictyostelium discoideum* is a good model system for exploring how the behaviors of individuals impart collective phenotypes and how social group performance is impacted by genetic and geographic relationships over a large spatial scale. These organisms exist as single-celled amoebae in the soil, but upon sensing imminent starvation, they coalesce into a multicellular organism (Kessin 2001). The amoebae signal to one another and aggregate over a centimeter or more to form a mound of ∼10^5^ cells (Kessin 2003). Over a period of several hours, the cells in the mound adopt an initial cell fate and form a migratory multicellular slug. The slug is attracted to sources of light and heat, which is likely to be an adaptation that directs it up to the surface of the soil. There, the slug transforms into a stalked fruiting body, consisting of a sorus (a ball of spores) that is lifted several millimeters above the surface of the soil by a thin stalk made of dead cells. The death of the stalk cells is altruistic, as their death likely confers benefits to the spores by lifting them up above the surface, enhancing dispersal or protecting them from dangers in the soil.

*Dictyostelium*’s unusual form of multicellularity is akin to a small society: amoebae of different genotypes can co-aggregate, resulting in slugs and fruiting bodies that are chimeric—that is, composed of more than one genotype (Fig. 1A). Like other societies where relatedness is below unity, *Dictyostelium* cell groups are also prone to conflicts of interest. Genetic diversity within the slug and fruiting body, combined with strong fitness consequences of having adopted the spore versus stalk cell fate (i.e., survival versus death), means that selection can favor selfish behaviors. In *Dictyostelium*, a genotype that can avoid the stalk fate or induce its partner to form it should have a selective advantage, referred to as cheating. Apparent cheating behaviors are indeed observed among different natural isolates when they are co-developed to form chimeric fruiting bodies (Strassmann et al. 2000; Fortunato et al. 2003; Buttery et al. 2009).

**Figure 1.**
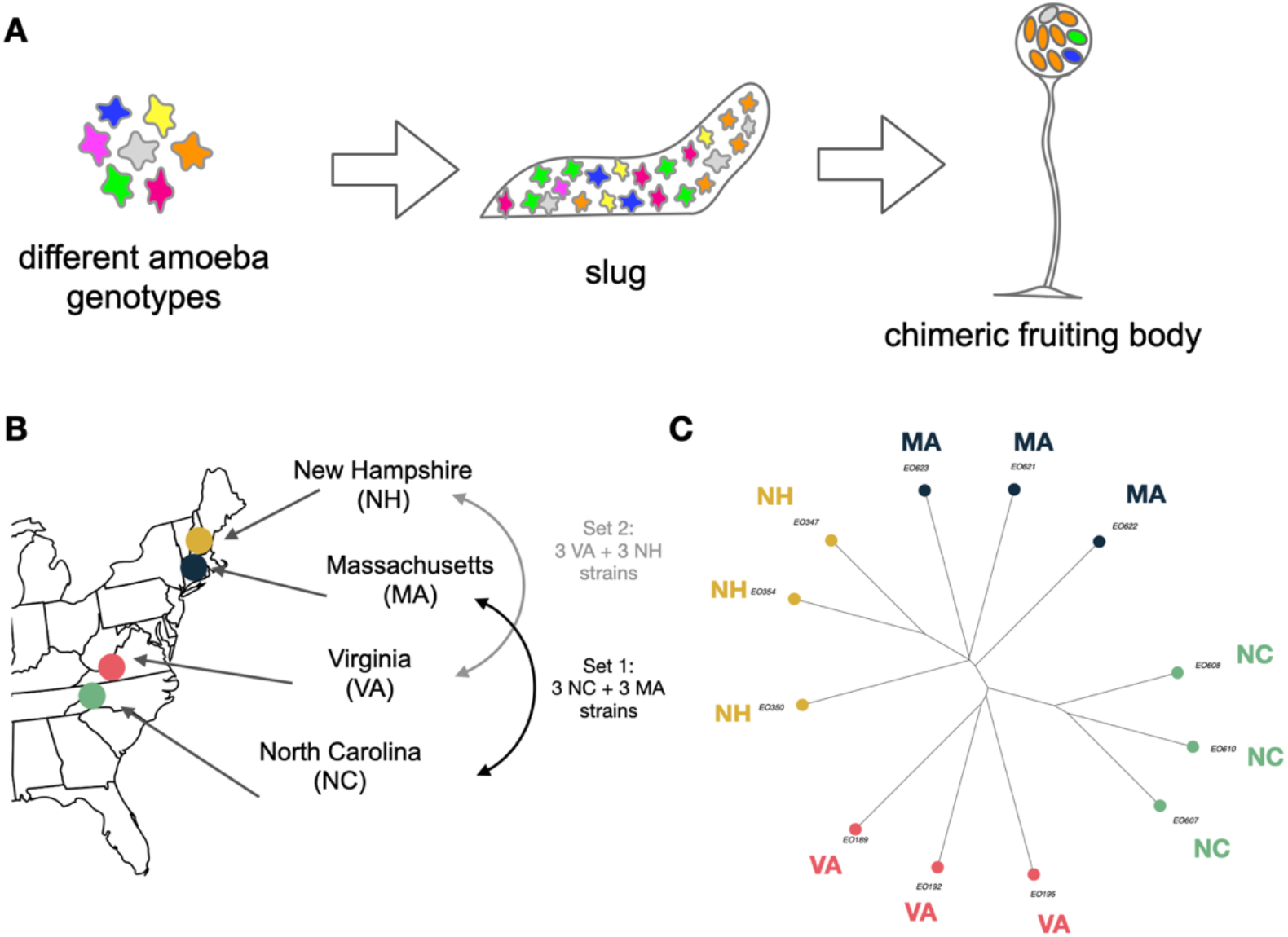
Chimerism in *Dictyostelium* as a model system to explore how genetic composition determines collective phenotypes. **(A)** When social amoebae are starved, the cells aggregate to form a multicellular organism in two stages: a migratory slug, followed by a fruiting body. Multiple strains (indicated by different colors) can co-aggregate to form a fruiting body, and it is possible to combine strains in many different combinations. In this work, the number and identity of strains that co-aggregate was varied while keeping the total number of cells constant. **(B) Experimental design**. Co-development assays were carried out for all possible combinations of two sets of six strains, resulting in 126 (= 63 × 2) unique combinations. Strains were collected from 10 × 10 cm plots at each of four locations: New Hampshire (NH), Massachusetts (MA), Virginia (VA), and North Carolina (NC). **(C) Genetic relationships among strains**. A distance-based tree showing overall genetic similarity among the 12 strains based on genome-wide SNPs.

To date, studies of chimerism in *Dictyostelium* have largely focused on whether spore production in chimeras is equitable—that is, on quantifying the extent of cheating. By comparison, less effort has been made to understand the consequences that chimerism entails for group-level phenotypes (although see Foster et al. (2002) and Votaw and Ostrowski (2017)). Yet, individual-level behaviors have the potential to impact the phenotype of the collective. For example, groups comprised of stalk-avoiding cells might produce fruiting bodies with shorter stalks, leading to poor spore survival. In short, group-level phenotypes have the potential to impact the cost-to-benefit ratio of the collective behavior and feedback to influence the decisions of individuals about whether to participate.

Indeed, perhaps owing to a history of conflict, strains of *D. discoideum* have evolved mechanisms that allow them to recognize and avoid unrelated strains following aggregation (Ostrowski et al. 2008). This kin recognition system involves a pair of heterophilic cell membrane proteins, *tgrB1* and *tgrC1*, that are highly polymorphic in natural populations (Benabentos et al. 2009). It is thought that these cell-surface proteins bind one another *in trans* across neighboring cells and that a sufficient match between the TgrB1 protein on one cell and the TgrC1 protein on another cell is required to proceed together through multicellular development. In the absence of a sufficient match, the cells partially sort out into different mounds that give rise to slugs and fruiting bodies (Benabentos et al. 2009; Hirose et al. 2011). Thus, *tgrB1* and *tgrC1* act as a “relatedness checkpoint” for proceeding with group formation and increase the relatedness among the cells in the fruiting body. Nevertheless, segregation of different genotypes is imperfect, and substantial chimerism usually persists into the fruiting body stage, at least under laboratory conditions (Ostrowski et al. 2008).

Here we take advantage of the *Dictyostelium discoideum* model system to quantify how group composition—specifically, the number of strains and their geographic and genetic relationships to one another—impacts the size, number, and productivity of collectives. We employ a statistical approach to detect and quantify genotype interactions involving the estimation of Walsh coefficients (Beerenwinkel et al. 2007; Weinreich et al. 2013; Poelwijk et al. 2016). This approach has been used previously to quantify genetic interactions among alleles that influence the phenotype of a cell, termed gene-by-gene interaction, or *g* × *g* (Sailer and Harms 2017b; Hall et al. 2019). Here we apply the same methodology to the conceptually analogous scenario of interacting genotypes and their effect on the phenotype of the group, termed genotype-by-genotype interactions, or *G* × *G* (Wade 2000; Wolf 2000).

In addition to quantifying the *G* × *G* interactions, we assess other metrics of group composition and their influence on collective phenotypes. We ask how the cooperative sporulation and morphology of fruiting bodies are altered by the number of strains in the group and their geographic and genetic relationships to one another. Specifically, we compare the sporulation of groups of strains from the same versus different soil samples, and we account for the degree of genetic divergence among strains in their ability to co-sporulate (Figs. 1B and C). Finally, we test the DEE and NNE hypotheses about whether and how *Dictyostelium* strains might be co-adapted (Fig. 2). The existence of either phenomenon can provide insight into the extent to which collective phenotypes have been shaped by a history of mutualistic cooperation or antagonistic conflict, respectively.

**Figure 2.**
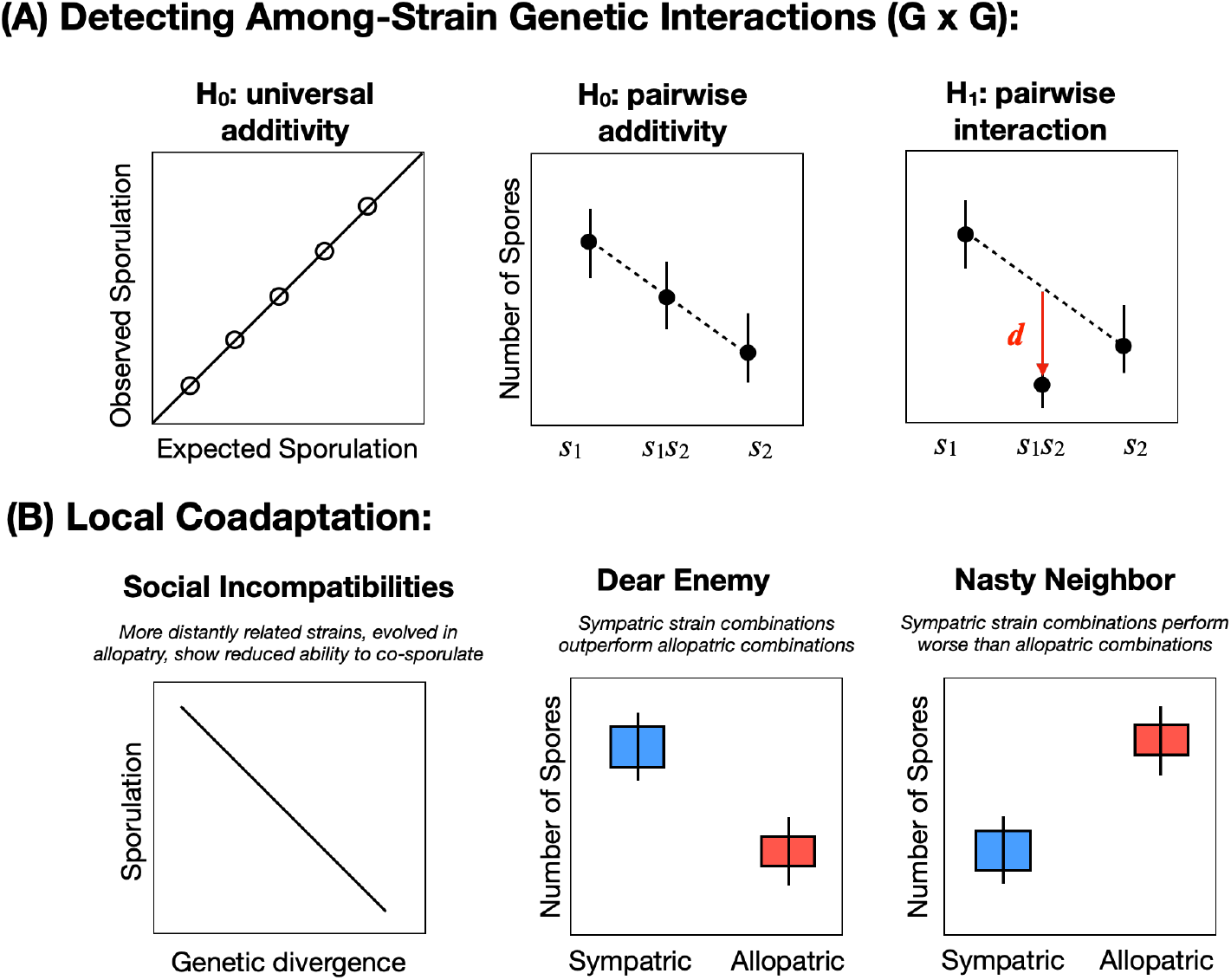
Experimental Predictions. **(A)** To test for evidence of genotype interactions (*G*×*G*), we compared the observed sporulation of strains developed in multi-strain mixes to an expected value based on the mean sporulation of the component strains when developed clonally. The deviation-from-expected sporulation (*d*) was calculated as this difference, divided by the expected sporulation. We assessed the evidence for pairwise and higher-order genotype interactions by estimating Walsh coefficients (see Methods and SI.) **(B)** We assessed whether the co-occurrence of strains in the same (=sympatric) vs different (=allopatric) soil samples led to higher or lower co-sporulation when co-developed together. Whereas higher sporulation of co-occurring strains could indicate mutually beneficial co-adaptation, worse sporulation of co-occurring strains could indicate enhanced levels of conflict owing to local competition. Finally, genetic divergence in allopatry could lead to incompatibilities that reduce the ability to co-sporulate. We tested this hypothesis by asking whether allopatric groups showed a significant negative correlation between among-strain genetic diversity and spore production.

## Methods

### Strain isolation

Table S1 provides a list of the strains and the GPS coordinates of the sites from which they were isolated. Briefly, three *D. discoideum* isolates were obtained from soil samples at each of the four sites shown in Figure 1B. Soil samples were collected from the surface of 10-by-10 cm plots in forested areas of Great Smoky National Park in North Carolina (NC), Smith MacLeish Field Station in Massachusetts (MA), Mountain Lake Biological Station in Virginia (VA), and Proctor Academy in New Hampshire (NH).

To isolate *D. discoideum*, 6 g of each soil sample was mixed with 30 ml of sterile water. The soil-water slurry was agitated briefly, and approximately 200 µl was deposited on hay infusion agar plates, with 400 µl of the bacterium *Klebsiella pneumoniae* as a food source. Spores were collected from a single fruiting body per plate and frozen at −80º C in KK2 buffer (per L: 2.25 g KH_2_PO_4_, 0.67 g K_2_HPO_4_) with 20% glycerol. Each isolate was subsequently cloned to ensure genetic homogeneity by plating the spores at low density on a lawn of *K. pneumoniae* on a Sussman’s medium plate (SM; Formedium, Ltd; with 2% agar) and picking fruiting bodies from the center of a single, isolated plaque. Spores of cloned isolates were grown to high density and then stored in KK2 buffer with 20% glycerol in a −80º C freezer.

### Overview of Experimental Design

We grouped the 12 strains into two sets of six, where each set contains three strains from two sites, as shown in Figure 1B. Set 1 consisted of three strains from North Carolina and three from Massachusetts (NC/MA). Set 2 consisted of three strains from Virginia and three from New Hampshire (VA/NH). Each strain set was used in co-development assays that consisted of all 63 possible combinations of the six strains. These combinations comprise 6 one-way (clonal), 15 two-way, 20 three-way, 15 four-way, and 6 five-way mixes, as well as one six-way mix. Co-development assays were carried out in a complete block design where all 63 mixes were assembled and developed on the same day. Each complete block was then replicated three to six times.

### Co-Development Assays

Each block of the co-development assays commenced by inoculating spores from the relevant frozen stocks onto lawns of *K. pneumoniae* on SM agar plates. Once fruiting bodies appeared, 5 × 10^5^ spores were collected from each strain, mixed with 400 µl of *K. pneumoniae*, and spread over SM plates. After two days, the cells were harvested from the SM plates during exponential growth. The cells were washed three times (340 *g* for 4 mins at 4°C) with cold KK2 buffer and resuspended at a density of 10^8^ cells/ml. Mixes were created by combining 60 µl of each of the required strains and then depositing a 100-µl aliquot of that mix (equal to 10^7^ cells) in a 4-by-4 square of a 47 mm gridded 0.45 μm nitrocellulose filter. For one-way mixes (i.e., single-strain controls), 100 µl of the cells were deposited on the nitrocellulose filter. The starting number of cells was thus the same for all combinations. The nitrocellulose filters were placed in 6 cm Petri dishes on top of Pall filter pads moistened with 1.5 ml of PDF buffer. The plates were incubated at 22°C with overhead light for two days.

### Sporulation Efficiency, Fruiting Body Size, Height, and Number

Once fruiting bodies had formed, the filters were imaged from the top and side before being transferred to 50-ml Falcon tubes containing two ml of detergent (490 mL KK2, 10 mL 0.5 M EDTA, 50 μl Igepal) and vortexed. Aliquots were counted on a Countess cell counter (ThermoFisher) to determine the number of spores. Sporulation efficiency was calculated as the number of spores divided by the number of starting cells.

The sorus diameter and the number of fruiting bodies was assessed by image analysis based on photos of the filters taken from above, whereas fruiting body height was determined by image analysis based on photos of the filters taken from the side. Side-images were cropped along the bottom, at the base of the fruiting bodies, and the midpoint of each sorus visible from the side was identified. The height of the fruiting body was then estimated as the distance from the middle of each sorus to the bottom of the cropped image. Image analysis was carried out in Image J. Example images and further details of the image analysis procedures are provided in the supplement.

### Genome Sequencing and Genetic Relationships Among Strains

Detailed methods for extraction of genomic DNA, library preparation, Illumina sequencing, and SNP filtering parameters are provided in the supplement. Briefly, we extracted genomic DNA from heat-shocked spores using a modified version of the protocol described by Adley et al. (2006). We used the Nexterra XT kit for Illumina library preparation, modified for quarter-sized reactions as described by Baym et al. (2015). We sequenced the library on an Illumina HiSeq X Ten sequencer. Reads were mapped to the *D. discoideum* reference genome using bwa-mem. SNPs were called using custom hard-filters in gatk using VariantFiltration with --filter-expression “QD < 2.0”, “FS > 60.0”, “MQ < 40.0”, “SOR > 3.0.” and –genotype-filter-expression “isHomVar == 1 && DP < 4.0”. Only biallelic SNPs without missing data were retained. The resulting VCF files were used to estimate Euclidean genetic distances among strains using the poppr package in R. A distance-based tree was constructed using the ‘ape’ package in R (Fig. 1C). For each mix, the average genetic distance was calculated by determining all possible pairwise combinations of strains and averaging their pairwise genetic distances (e.g., for mix 123, the average of the nine combinations of pairwise encounters—11,12,13,21,22,23,31,32,33).

### Statistical Analyses

The deviation from expected sporulation was calculated as the difference between the observed and expected sporulation efficiency of a mix, divided by its expected sporulation: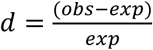, where *obs* is the observed sporulation efficiency of the mix, and *exp* is the expected value. The expected sporulation efficiency was calculated as the arithmetic mean of the sporulation efficiencies of each strain when developed as a monoculture (i.e., clonally). Negative values of *d* thus indicate that the group performed worse than expected, whereas positive values indicate the group produced more spores than expected, based on the strains’ clonal sporulation.

We assessed overall evidence for additivity by regressing the observed sporulation of a group on its expected sporulation. Slopes and intercepts were estimated using standardized major axis regressions to account for error in both the independent and dependent variables. Regressions were estimated in R using the package ‘smatr’, with slope.test=1 and elev.test=0. Principal components analysis was carried out using the ‘prcomp’ function in R, with centering and scaling. Statistical models of observed sporulation as a function of expected sporulation, geography (allopatry/sympatry), number of strains, genetic distance, and block were carried out in R using glm. Owing to the small number of sites and strains, all factors were modeled as fixed effects.

### Epistasis Coefficients

To estimate first-order (i.e., individual strain), pairwise, and higher-order epistasis coefficients, we use the group sporulation data to calculate Walsh coefficients (Weinreich et al. 2013; Sailer and Harms 2017a). Estimation of Walsh coefficients involving mutation effects typically includes the phenotype of a wild-type genotype, which harbors no mutations. There is no wild-type counterpart in our experimental design because sporulation efficiency can only be measured when at least one strain is present. Instead, we used the grand mean sporulation efficiency of all mixes in the relevant strain set as an estimate of the baseline sporulation efficiency. To estimate the relative importance of first-, second-, and higher-order interactions on group sporulation, we calculated the correlation between the observed and estimated sporulation of mixes in each strain set, where estimated sporulation accounted for the tested and lower-order interactions (Sailer and Harms 2017a). This analysis provides an estimate of the proportion of overall variation in sporulation of the group that can be explained by the sporulation efficiency of single strains or pairwise interactions between them. Confidence intervals on these estimates were estimated by bootstrapping over strain combinations of a particular interaction order. For example, for each of the two geographically distinct strain sets considered, the effect of pairwise interactions was determined by bootstrapping over the 15 two-strain combinations. Further explanation of the estimation of Walsh coefficients is provided in the supplement.

## Results

### Group phenotypes depend on non-additive social interactions

To better understand the nature and effects of interactions between individuals on *D. discoideum* group phenotypes, we first asked whether there was a significant relationship between the expected and observed sporulation of groups. Expected sporulation was calculated as the average of sporulation of the component strains when developed in single-strain monocultures. Under the null hypothesis of complete additivity, we expected to see a slope of 1 and an intercept of 0. For NC/MA strains, however, this null hypothesis was rejected (slope estimate = −1.86, intercept= 3.18, *P*<0.0001; Fig. 3A). In addition, there was no significant relationship between observed and expected sporulation (*r* = −0.07, *df* = 55, *P*=0.59). For VA and NH strain combinations, the null hypothesis of a slope = 1 and an intercept = 0 was also rejected (Fig. 3B; slope estimate = 2.48, intercept = −1.58, *P*<0.0001). However, the relationship between observed and expected sporulation was significant (*r*=0.34, *df* = 55, *P* = 0.01). These results indicate that sporulation patterns deviate significantly from what would be expected if sporulation were purely an additive function of the sporulation of the group’s component strains.

**Figure 3.**
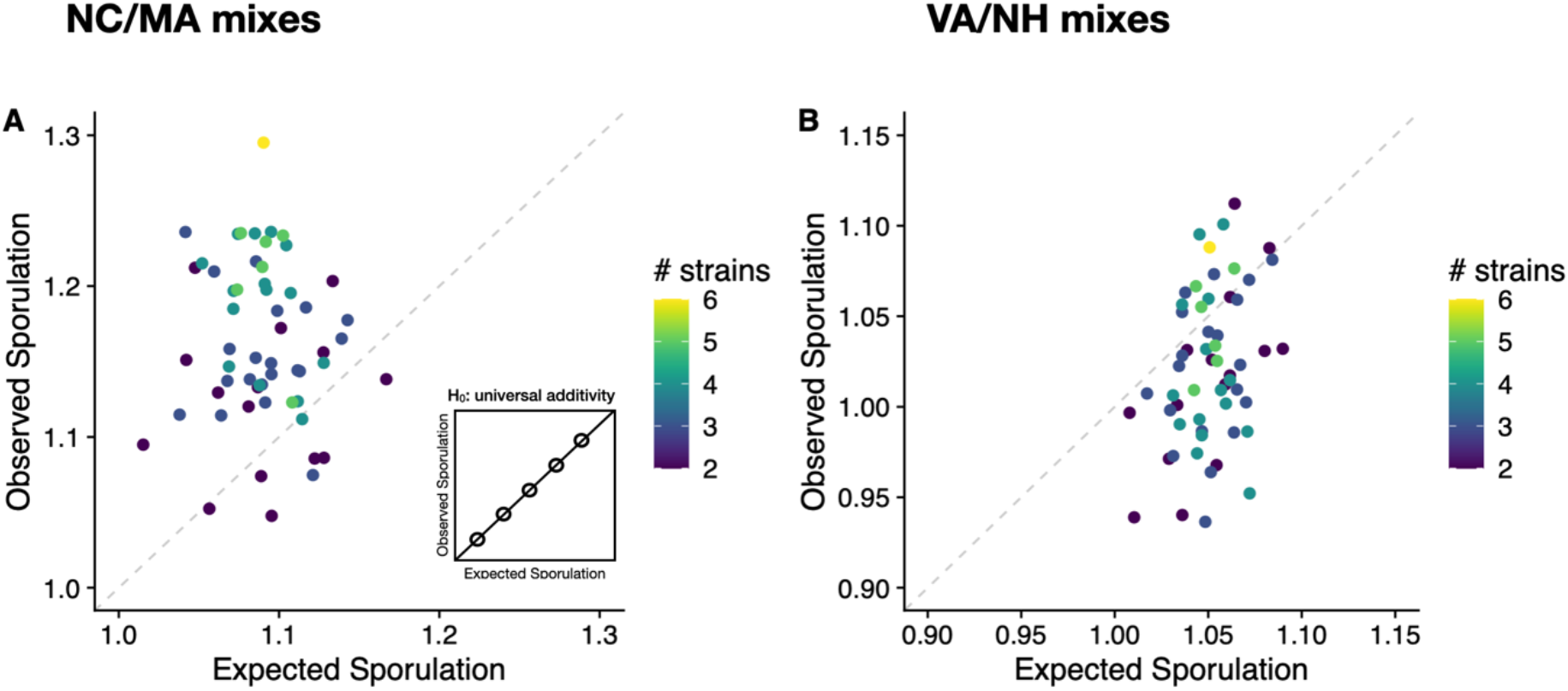
Overall sporulation pattern shows only weak additivity. If spore production is an additive function of the component strains, then group sporulation should be predictable from the sporulation efficiencies (the number of spores, divided by the number of starting cells) of each strain developed clonally (i.e., ‘expected sporulation’); see inset. Each point represents a unique strain mix, and colors indicate the total number of strains in the mix. **(A)** NC/MA strain combinations. **(B)** VA/NH strain combinations. See text for details.

The mean effect of mixing on sporulation efficiency also differed between the two sets of strains. For NC/MA strain combinations, the mean deviation from expected sporulation was significant and positive (=7%), indicating that combining strains tend to enhance the spore production of the group (one-sample *t-*test: *t*=8.99, *df*=56, *P*<0.0001). This finding was largely driven by the NC strains, which produced many more spores as a group than expected based on their sporulation in monocultures. In contrast, the MA strains performed somewhat worse than expected. For VA/NH strain combinations, the average deviation between expected and observed spore production was significant and negative (−3%; one sample *t-*test: *t* = −5.2, *df*=56, *P*<0.0001), indicating a cost of mixed groups on spore production, at least with respect to spore number. We discuss these results in more detail in later sections.

### Pairwise and higher-order genotype interactions influence group sporulation

To determine the nature of interactions leading to group effects on spore production, we estimated Walsh epistasis coefficients for the two groups of strains (NC/MA and VA/NH). These coefficients quantify strain interactions as the deviation of each strain combination from its expected sporulation across all mixes in which it occurs. Walsh coefficients for both sets of mixes (NC/MA and VA/NH) revealed a variety of individual interactions (Fig. 4). These results also show that the main effects (i.e., additive effects of single strains), pairwise, and higher-order interactions all contribute significantly to the sporulation efficiency of mixes (see Fig. 4 insets). Of note, the combined contribution of interaction effects was similar in magnitude to that of the additive effects. This result indicates that, while each strain can independently influence the sporulation efficiency of its group, interactions among strains also have a substantial impact on group outcomes.

**Figure 4.**
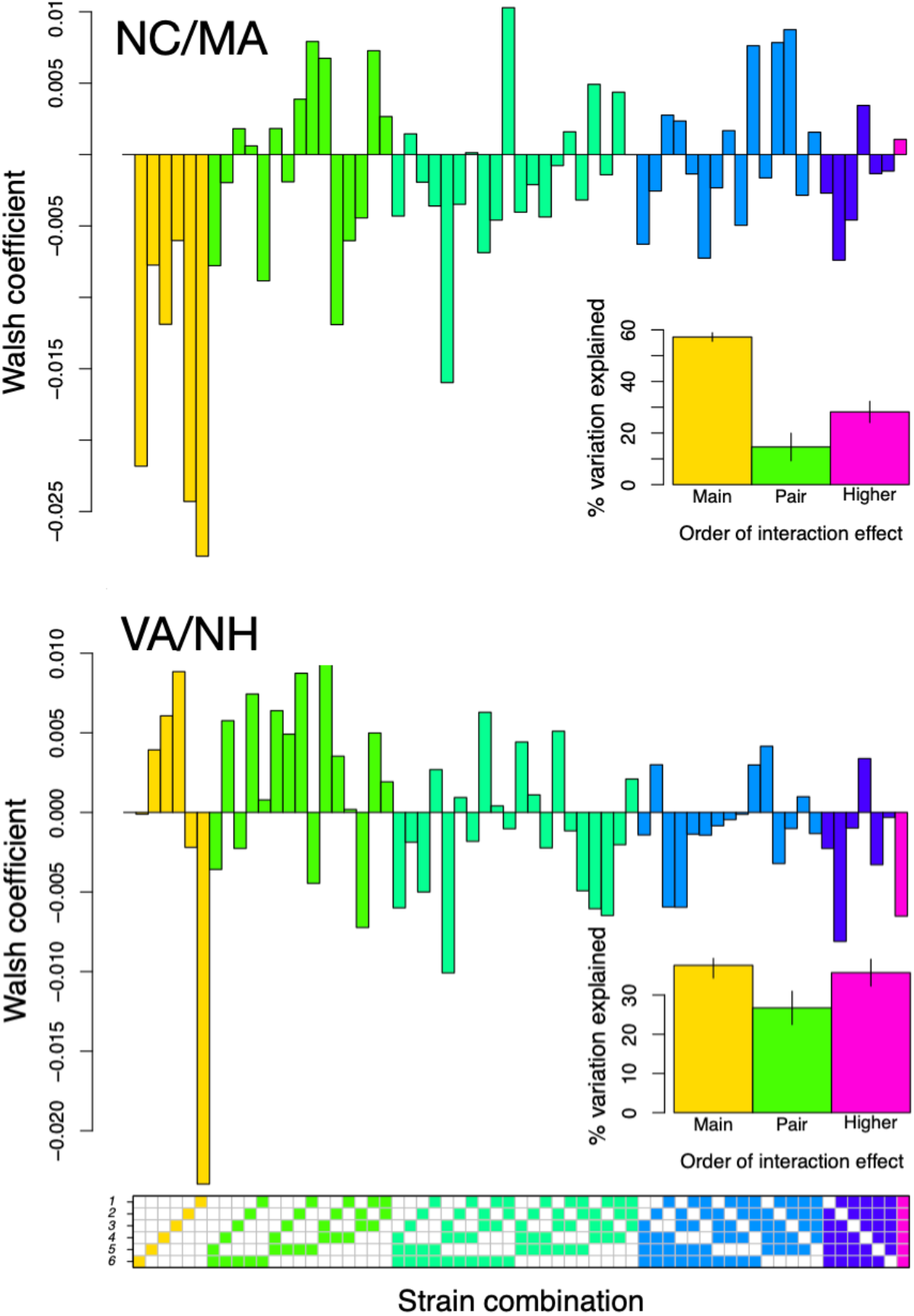
Estimates of Walsh coefficients. The matrix beneath the figure indicates the set of strains (1-6; see Table S1) mixed and analyzed to estimate the Walsh coefficient directly above. Each column of the matrix indicates one of the 63 possible mixes, and each row indicates the presence or absence of each strain in that mix, with shading of the cell indicating that the strain was present. Yellow: single strain (i.e., clonal) development. Light green, green, light blue, dark blue, and magenta represent the 2-way, 3-way, 4-way, 5-way, and 6-way mixes, respectively. Inset: percentage of variation in sporulation that can be explained by the main effects of each strain versus pairwise or higher-order interactions among strains. Error bars are calculated based on bootstrapping.

### Some strains have strong and consistent effects on their groups

Prior work has shown that collective phenotypes can be influenced by the presence of certain keystone individuals that drive the behavior of their groups (Sih and Watters 2005; Sih et al. 2009; Modlmeier et al. 2014). We assessed whether some strains might act as ‘bad apples’ that consistently lower the group’s productivity, below what is expected based on how well members sporulate individually. We also looked for the converse— strains that consistently elevate the sporulation of their groups—which we refer to as ‘good eggs’. To do so, we used ANCOVA to determine how the presence or absence of each strain impacts the sporulation of its groups after controlling for their expected sporulation (Fig. S3). These analyses show that several strains are ‘good eggs’: strains 1, 2 (both NC), strain 6 (from MA), and strain 1 (from VA). Adding these strains to a group increases its sporulation by ∼4% from its expected values. Our analyses show a consistent negative effect for only one strain, strain 4 (from NH); groups with this strain do 2.8% worse than expected based on their clonal sporulation. Together, these results indicate that certain strains interact with the presence of other strains in a consistent way.

### Strain diversity enhances sporulation and alters fruiting body morphology

To address the sporulation patterns we observed, we also asked whether increasing the number of strains in a mix enhances or reduces the sporulation of the group. We expected that sporulation might decline as the number of strains in the group increases, in part because many of these strains are genetically and geographically distant (>1000 km) from one another (Fig. 2). We hypothesized that lack of interaction among populations over their evolutionary history could lead to divergent evolutionary trajectories and generate incompatibilities that would be evident in mixes of allopatric strains. Moreover, increasing the number of strains in a mix might increase the probability of sampling at least one incompatible pairing.

Given these expectations, we were surprised to see the opposite relationship (Fig. 5). For NC and MA strain combinations, there was a significant positive correlation between the number of strains in the mix and the number of spores they produced. For VA/NH strains, the relationship was not significant, but the sign was also positive, and the 5-way and 6-way mixes were amongst the highest performing mixes in terms of spore production. At least for this metric of productivity, increased strain diversity was associated with improvements in group performance, suggesting that *D. discoideum* shows social heterosis.

**Figure 5.**
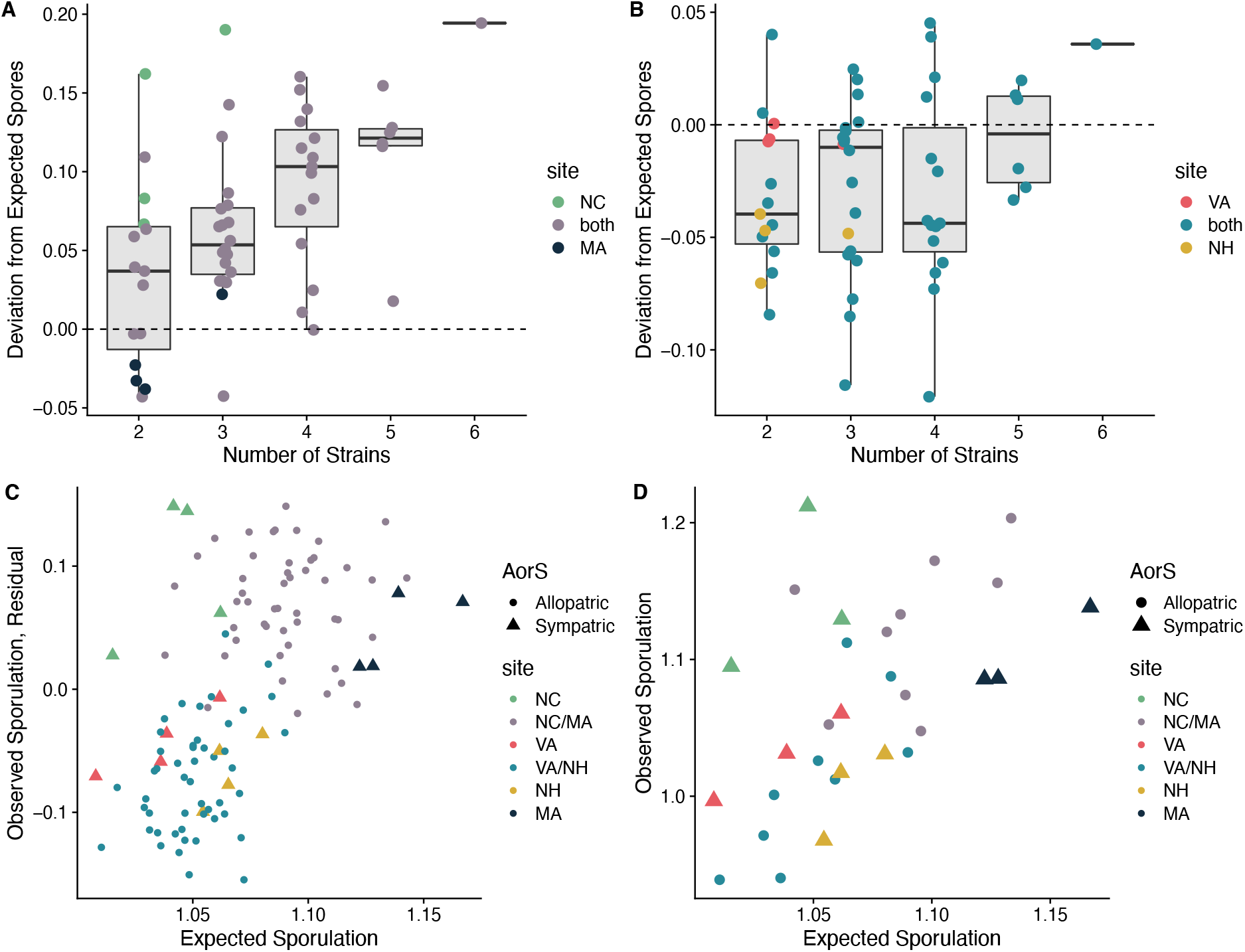
Genetically diverse groups sporulate as well as, or better than, clonal groups. **(A)** NC/MA strain combinations or **(B)** VA/NH strain combinations. Each point represents a unique mix of strains, and colors indicate the geographic site where the strains were isolated. A deviation of zero (indicated by the dotted line) indicates the mean sporulation of the six strains when developed clonally. Positive and negative values of the deviation indicate that the mix produced more or fewer spores, respectively, than expected based on the sporulation of the component strains when developed clonally. In both sets of mixes, the sporulation of the 5-way and 6-way mixes is as high—or higher than—the 1-way (clonal) development. **(C)** Controlling for the effect of strain number (i.e., plotting the residuals from a model that contains only the number of strains as an explanatory factor) or **(D)** limiting the analysis to two-way mixes shows that the relationship between observed and expected sporulation is linear (*F*_1,28_ = 12.3, *P* = 0.002).

Spore production of the group is one metric by which we can measure the success of fruiting body formation. However, additional group attributes might influence the collective’s success, aside from the number of spores they produce. For this reason, we also measured fruiting body size, number, and shape (Fig. 6). For NC/MA strain combinations, as the number of strains in the group increased, fruiting body size declined and the number of fruiting bodies increased. For VA/NH mixes, we also saw an initial decline in fruiting body size with increasing strain number, although it partially rebounded for the 5-way mixes and the 6-way mix. Thus, diverse groups produced more spores, but their fruiting bodies were smaller and more numerous than expected (Fig. 6B).

**Figure 6.**
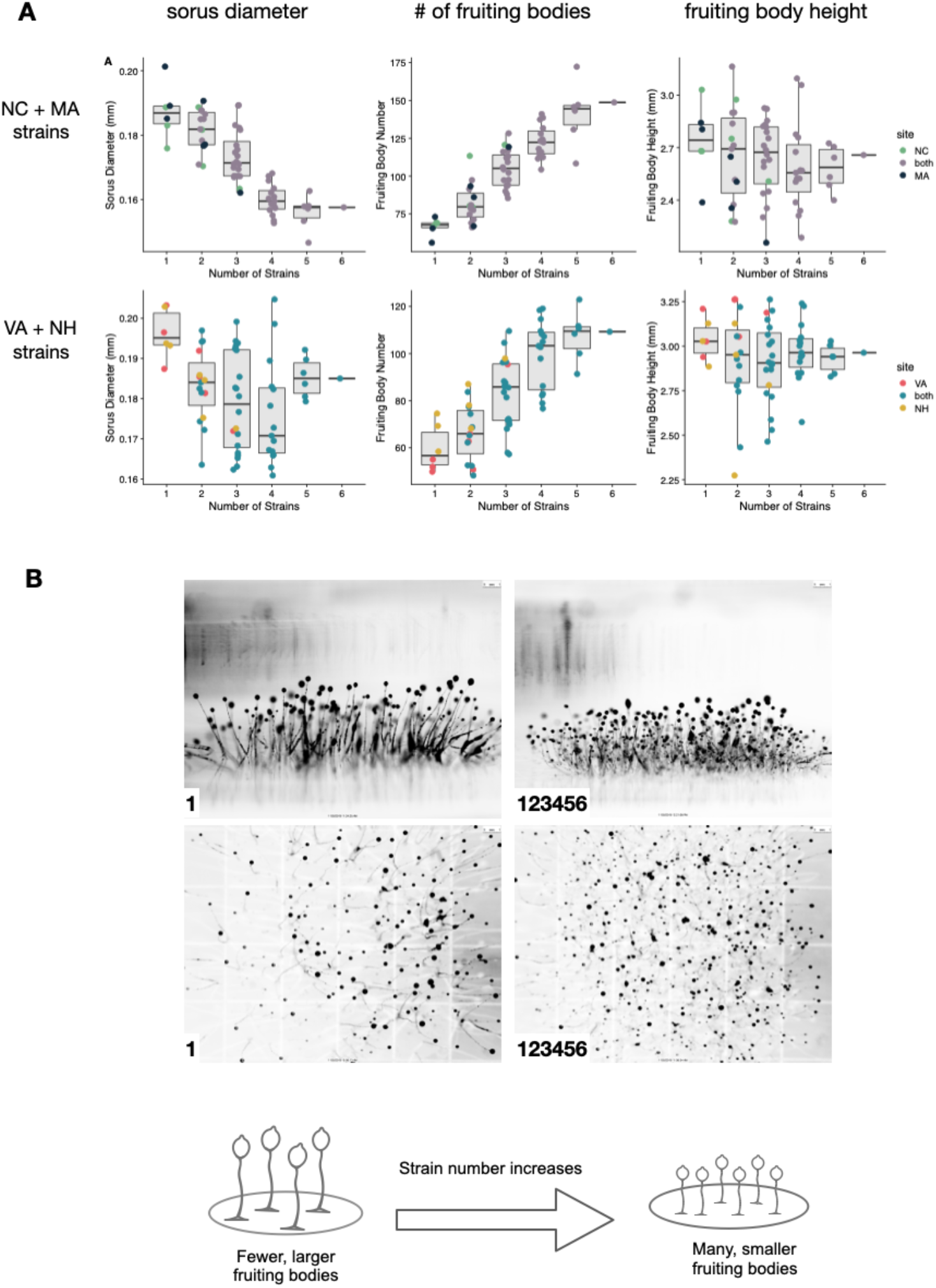
Genetic diversity in groups impacts fruiting body size, number, and shape. **(A)** Joint phenotypes: fruiting body size (sorus diameter), number of fruiting bodies, and fruiting body height as a function of the number of strains in a group. **(B)** Example images of fruiting bodies imaged from above or the side. The left image shows clonal fruiting bodies produced by strain 1, and right images shows fruiting bodies of the 6-way mix. Colors have been inverted and contrast adjusted for illustration purposes only. Additional images and the details of the automated image processing are provided in the supplement.

Given that the number of starting cells was held constant across all mixes and that little-to-no cell division takes place during the formation of the fruiting bodies (Chen et al. 2004), the most likely explanation for smaller fruiting bodies is that the cells divide themselves up into more groups, each of smaller size. Smaller fruiting bodies should also have shorter stalks, such that spores are not lifted as far off the ground. For fruiting body height, we see a slight trend towards shorter fruiting bodies, although we note that our side-images of fruiting bodies can provide only an approximate estimate of height (Fig. 6B and Fig. S2).

### Dear enemies or nasty neighbors? Do sympatric strains show enhanced conflict or synergism?

Within populations, selection might favor genotype combinations that work well together, particularly when the outcome of cooperation is a dispersal structure that promotes access to novel competitors (i.e., when cooperation is local, but competition is global). Alternatively, if neighbors are also each other’s closest competitors, then we could instead observe worse-than-expected sporulation in mixes of neighboring strains owing to enhanced conflict when fitness is a zero-sum game and benefits must be achieved at cost to one’s neighbors (Fig. 2B).

Geography had inconsistent effects on the ability of strains to co-sporulate (Fig. 7; Table S2). For example, pairs of NC strains sporulate significantly better with one another than expected (one-sample *t*-test: *t* = 4.19, *df* = 3, *P* = 0.02). However, for MA strains, mixing has a negative effect on average, although the deviation from expected is not significantly different from zero (one-sample *t*-test: *t* = −1.30, *df* = 3, *P* = 0.28). However, all three groups (NC, MA, and NC/MA) are significantly different from one another (*F*_2,165_=5.41, *P*=0.005; all pairwise *P*<0.04 based on t-tests with Holm’s correction for multiple comparisons). Thus, while MA strains do not co-sporulate significantly worse with one another than they do clonally, they are significantly worse together than NC strains are and also worse than allopatric (NC/MA) combinations, as the latter two groups showed positive deviations from expected sporulation, indicating that the chimeric treatments had produced more spores than expected.

**Figure 7.**
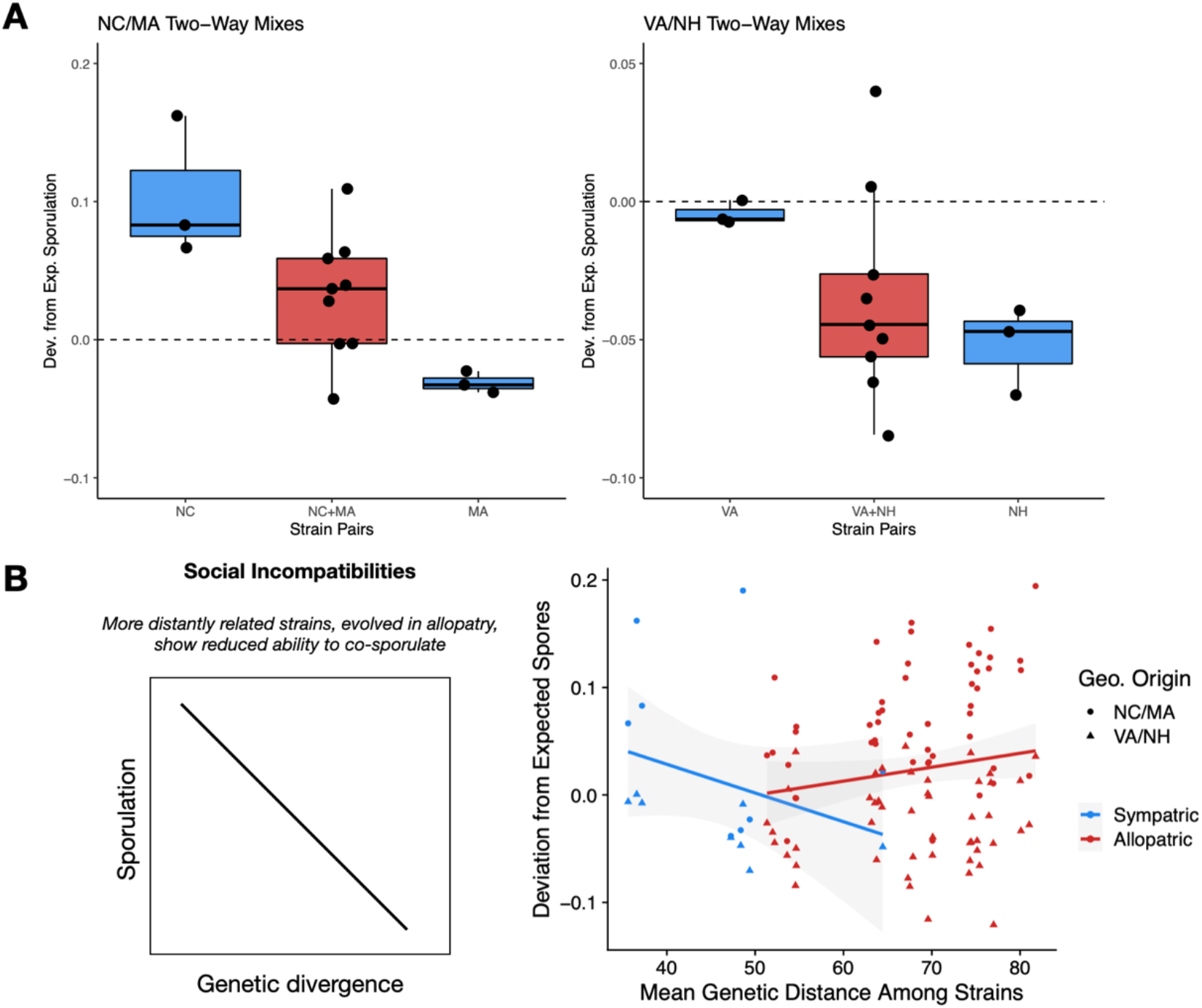
Tests of co-adaptation among sympatric strains and incompatibilities as a function of genetic or geographic distance. **(A)** Are co-occurring strains dear enemies or nasty neighbors? North Carolina (NC) strains co-sporulate significantly better than expected, whereas Massachusetts (MC) strains do significantly worse. Virginia strains (VA) co-sporulate nearly exactly as expected, whereas New Hampshire (NH) strains perform worse than expected. Shown here are pairwise (two-way) mixes only, which avoids the confounding effect that the number of strains has on sporulation. **(B)** No evidence of reductions in sporulation as a function of the genetic divergence between strains, particularly for allopatric strains, in contrast to our predictions.

VA strains did not co-sporulate significantly differently with one another than expected, based on their clonal sporulation (one-sample *t*-test; *t* = −2.66, df = 3, *P* = 0.08), but NH strain combinations did produce significantly fewer spores together than expected (one-sample *t*-test; *t* = −7.75, df = 3, *P* = 0.004), as did the VA/NH allopatric mixes (one-sample *t*-test; *t* = −4.54, df = 48, *P* < 0.0001). However, the differences among these groups were not significant (*F*_2,336_=1.64, *P*=0.196). Taken together, these results reveal a wide range of interactions among sympatric strains: from synergism and enhanced spore production as a group (NC strains) to performance that is significantly worse-than-expected (NH strains) and even worse than allopatric combinations (e.g., MA strains do worse with each other than they do with NC strains.)

### Limited evidence that evolution in allopatry leads to accumulation of social incompatibilities

A major hypothesis in social evolution is that social interactions among conspecifics lead to adaptations and counter-adaptations that drive region-specific co-evolutionary trajectories (West-Eberhard 1979, 1983). Divergent evolutionary trajectories might then generate social incompatibilities that prevent cooperation from taking place when allopatric individuals encounter one another. To test this possibility, we asked whether there is a reduction in sporulation, a cooperation-dependent phenotype, with increased genetic divergence among the strains.

We did not find that increased genetic divergence was associated with reductions in sporulation for allopatric mixes, defined as those where at least two strains come from different sites (Fig. 7B). The correlation was positive in sign and not significant (*r* = 0.16, *df* = 96, two-tailed *P* = 0.11). For sympatric strains, the relationship was negative and also not significant (*r* = −0.33, df = 14, *P* = 0.22). Although each correlation was not significantly different from zero, the two correlations were significantly different from one another (*P*=0.03, confidence intervals and difference estimated using ‘twocorci’ package in R).

### Putting it all together: how does the composition of groups influence collective phenotypes?

We used principal components analysis (PCA) to develop a better understanding of the relationship between the different group morphological attributes, sporulation, and how these traits are influenced by the number of strains in the group and the genetic differences between them (Fig. 8). Approximately 84% of the variance is explained by the first three principal components. The analysis confirms our overall picture of the results: as the number of strains increases, the number of fruiting bodies increases and their size (sorus diameter) decreases. In other words, they produce more, yet smaller fruiting bodies (Fig. 6B). In addition, the PCA shows that increases in spore number (and a positive deviation from expected sporulation) are inversely related to the fruiting body height. This result suggests that greater spore production is indeed associated with reduced stalk investment—and a potential cost in terms of lifting the spores above the substrate. Finally, while the genetic and geographic relationships between the component strains contribute strongly to PC2, the relationships of these explanatory factors to the morphological attributes are more orthogonal to the morphological variables, illustrated by the distinct rotation of these eigenvectors.

**Figure 8.**
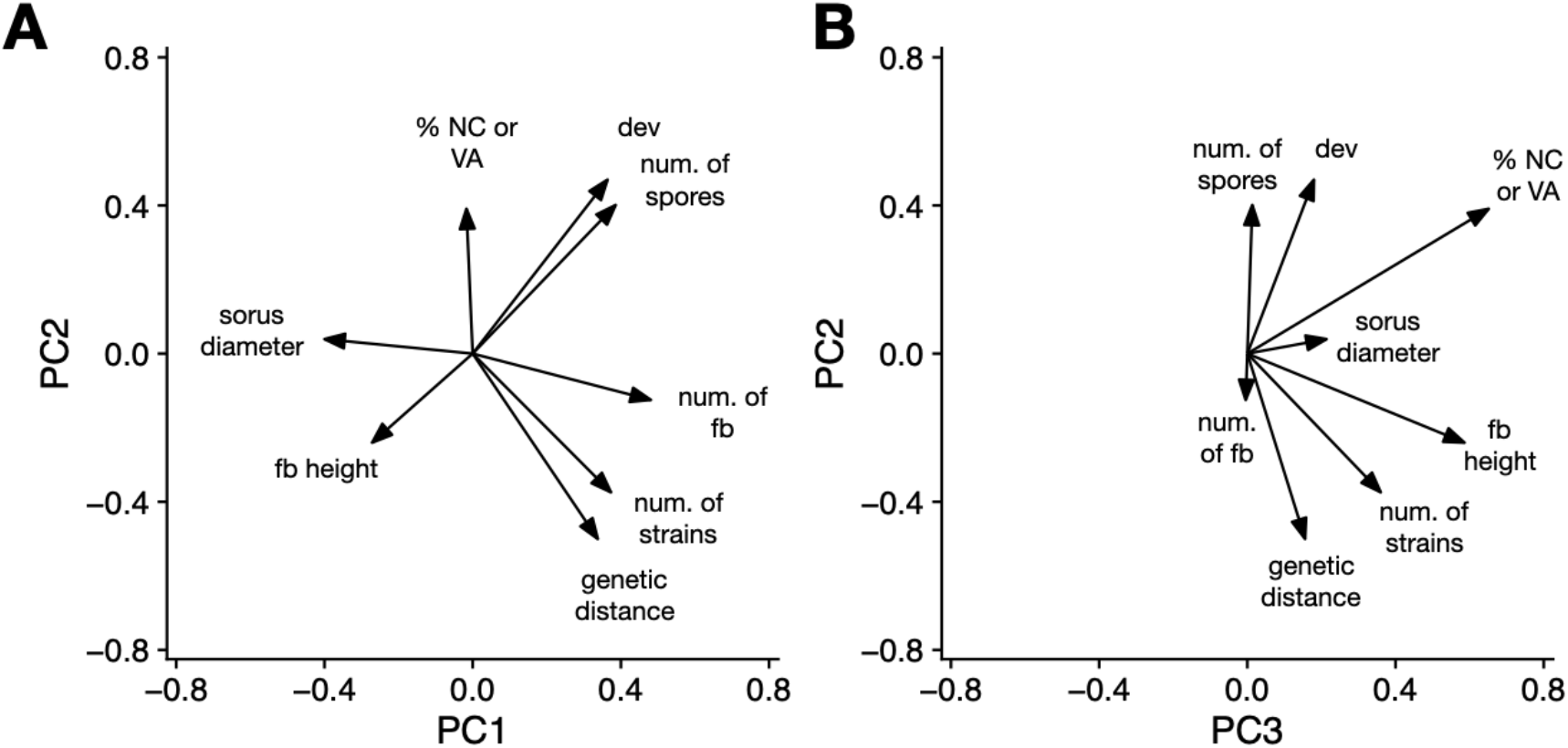
Principal components analysis shows the relationship between the number of strains in a mix, their genetic and geographic relationships, and the results in terms of group productivity (spore production) and morphological traits (fruiting body number and height and sorus diameter). (A) principal component (PC) two versus PC1, (B) PC2 vs PC3.

To assess the statistical significance of each explanatory factor, we modeled the observed sporulation of each group as a function of its expected sporulation (i.e., mean clonal sporulation, which accounts for any differences among strains in clonal spore-stalk allocation), geography (allopatry versus sympatry), genetic distance, and the number of strains. These analyses are summarized in Table S3. Consistent with the PCA results, we see strong effects of expected sporulation, indicating some predictability of the group as a function of the clonal behavior of the strains, and the number of strains in the group. Allopatry versus sympatry and genetic distance are both significant in this model, reflecting that these factors have some influence on sporulation, once other factors have been accounted for.

### Kin recognition might explain the morphological changes, but also the increased sporulation of genetically diverse groups

Our results showed that increases in strain number led to decreases in the size of multicellular fruiting bodies but increased total spore number. As strain number increases, cells of any given strain become increasingly surrounded by non-identical cells. For example, in the six-way mix, each strain is present at a frequency of 1/6, or 17%, and thus every genotype is in the minority. We hypothesized that both patterns (smaller groups yet more spores) might result from kin discrimination mediated by non-matching alleles of different strains at *tgrB1* and *tgrC1*. Kin recognition involves an initial stage where cells of multiple genotypes co-aggregate, followed by a process where different genotypes partially sort out, resulting in subdivision of the original group (see Ostrowski et al. (2008)). Therefore, it makes sense that recognition will have the side-effect of creating smaller groups. Moreover, in *Dictyostelium*, these groups may experience a cost in terms of spore dispersal owing to the smaller fruiting bodies that result and their commensurately shorter stalks.

In contrast to the predictable effects of genotypic diversity on group size, which makes sense given what is known about how kin recognition works in *Dictyostelium*, the observation that more diverse groups also produced more spores was more puzzling, and the most parsimonious explanation is that cells alter their spore-stalk allocation in response to detection of non-self. We thus wondered whether kin recognition has two distinct effects on social group formation in *Dictyostelium*: (1) avoidance of non-kin, leading to smaller groups and fruiting structures, with potential consequences for effective dispersal, and (2) increased propensity for selfishness in groups that do consist of non-kin by shifting of cell allocation towards the spore fate, potentially at a cost to the production of the stalk.

Our experiments do not address this question directly, but we took advantage of existing RNA-seq datasets to ask whether *tgrB1*-*tgrC1*-mediated recognition is associated with facultative shifts in the probability of altruistically becoming a stalk cell. Hirose et al. (2015) determined which genes show altered expression when cells are surrounded by non-self cells (i.e., strains with non-compatible *tgrB1*-*tgrC1* alleles) compared to control assays where cells are surrounded by genetically identical cells. They determined this change in gene expression at two timepoints during multicellular development: at 8-hr, coinciding with the initiation of *tgrB1*-*tgrC1* expression and signaling, and at 12-hr, just after cells have adopted an initial cell fate—referred to as ‘prespore’ or ‘prestalk’ cells (Benabentos et al. 2009; Hirose et al. 2015). We merged the genes identified in this study with a different dataset that classified genes according to their degree of prespore or prestalk cell expression bias (Parikh et al. 2009). If detection of non-self cells increases the probability that a cell adopts the prespore fate, then we expect to see an upregulation of prespore genes and/or a downregulation of prestalk genes. Analysis of these two datasets revealed that the genes upregulated in response to non-self are not, on average, prespore-biased (Fig. S4). However, the downregulated genes were strongly prestalk-biased (one-sided *t* = −4.98, *df* = 136, *P*<0.0001; Fig. S4). This result suggests that interaction with unrelated strains leads to either reduced expression of prestalk genes or fewer prestalk cells, and it supports our hypothesis that the increased sporulation of genetically diverse groups could be mediated by cellular responses acting downstream of *tgrB1-tgrC1*, such that those cells that remain in a chimeric structure following kin recognition also behave more selfishly.

## Discussion

We created collectives of different *Dictyostelium* strains to explore how the genetic composition of a group influences its collective phenotype. We dissected the genetic basis of a complex collective phenotype into components capturing the additive contribution of each strain to its group’s function, as well as the interactions between genotypes, known as *G* × *G* epistasis (Wade 2000; Wolf 2000; Lambrechts 2010). Our analyses thus allow the quantification of the relative contributions of these different types of genetic variance to joint phenotypes—here, spore production and fruiting body morphology—that are likely to be closely tied to fitness in nature. Our experimental design is novel in that it applies experimental approaches and mathematical analyses that have been successfully used to quantify the form and extent of *g* × *g* epistasis (that is, genetic interactions among alleles in an individual) to the detection of epistasis at a higher level of biological organization.

Our findings indicate a consistent influence of particular strains on group phenotypes, but also the existence of extensive pairwise and higher-order epistasis. One factor that strongly influenced the group morphology and behavior was the number of strains in the group. Although we had predicted that more strains (i.e., more genetic diversity) would lead to the production of fewer spores, we found that groups with more strains tended to have greater, rather than lesser, spore production. This effect was mostly driven by one set of strains (from North Carolina and Massachusetts). However, even in the second set (from Virginia and New Hampshire), sporulation fully recovered to expected levels in the five-way and six-way mixes. Thus, high levels of genetic diversity were not associated with major declines in sporulation. At a minimum, this result indicates that severe incompatibilities have not emerged across allopatric populations and that *Dictyostelium* retains a high ability to cooperate with distantly related strains.

The above finding is reminiscent of the unicoloniality that is observed in some ant species that cooperate with other groups across their geographic range, despite low relatedness (Helanterä et al. 2009). How and why unicoloniality evolves remains a mystery, but it might reflect ecological or demographic changes (e.g., population expansions into new territories) that influence the cost and benefits of cooperation and allow unrelated groups to benefit from ecological dominance. Either way, *Dictyostelium*, which itself has likely dispersed from a glacial refugium, may hold keys to this unusual phenomenon, which has been suggested to be the next ‘major transition’ in complexity (Bourke 2011).

While we did not observe major costs of strain diversity on spore production, there were morphological changes associated with increases in diversity of the mix, and we hypothesize that these changes may have important impacts on selection in nature. In diverse groups, the diameter of the sorus (the spore-bearing head of the fruiting body) was smaller and the stalk declined slightly in height, while the total number of fruiting bodies increased. This result shows that the same number of starting cells divided into more groups, producing more yet smaller fruiting bodies. Several lines of evidence suggest that lifting spores off the ground is an important and strongly selected trait—for example, stalked fruiting bodies have independently evolved multiple times across the eukaryotic phylogeny, suggesting a common fitness benefit to lifting spores off the substrate (reviewed in Broersma and Ostrowski 2022). The collective phenotype of the group is thus modified by the interactions among diverse strains in ways that have the potential to influence fitness negatively.

To date, studies with *Dictyostelium* have focused primarily on one type of fitness cost that cells might experience when they join forces to build a collective: they might be cheated on, whereby one genotype contributes more than its fair share to the production of the dead stalk, thus gaining the benefits of the stalk without having to pay the full cost (Strassmann et al. 2000; Strassmann and Queller 2011). Cheating has the potential to be strongly selected, and so many studies have focused on quantifying the fairness of spore production. By comparison, less attention has been paid to how chimerism influences the group morphology and the benefits of partaking in the collective behavior in the first place (but see Foster et al. 2002; Votaw and Ostrowski 2017). Our results point to costs of chimerism that emerge via the group’s collective phenotype—specifically, the size, height, and the number of fruiting bodies produced. In addition, while costs of chimerism might select for avoidance of genetically different strains, in this case, kin recognition itself is its likely cause. This finding demonstrates the logic behind Crozier’s famous paradox for why kin recognition is not the answer to cheating—those who are choosy will not gain the full benefits of cooperation (Crozier 1986). Indeed, as others have pointed out, *Dictyostelium*’s highly imperfect kin recognition system may be more a feature than a bug, and the benefits of greater group size can potentially outweigh the risks of being cheated (Ostrowski 2020; Pentz et al. 2020; Márquez-Zacarías et al. 2021).

Our results show mixed support for the dear enemy and nasty neighbor phenomena. For example, mixes of NC strains show significant elevations in their sporulation, potentially indicative of a dear enemy effect. Conversely, NH and MA strains co-sporulate poorly with strains in their immediate vicinity, a pattern more indicative of nasty neighbors. However, the bad apple/good egg analysis modifies our interpretation of this result. Two of the three NC strains were classified as good eggs, meaning they increase the sporulation of groups they are added to—but, by definition, they do so regardless of whether those groups consist of sympatric or allopatric strains. Similarly, while NH strains do significantly worse together than expected, our only example of a bad apple was strain 4 from NH. This strain brings down the sporulation of groups it was added to, and it does so regardless of the geographic origin of its partner strains. Thus, while sympatric strains can do somewhat better or worse together than allopatric strains, these effects may be better explained by the fact that some sites harbored more good egg or bad apple strains than others. In other words, the presence or absence of key behavioral types may play a bigger role in the group characteristics than co-evolution between them (Modlmeier et al. 2014; McAuliffe et al. 2015; Wheatcroft and Price 2018). Similarly, recent work has demonstrated that interactions between specific mutations can drive general trends in mutation effects (Lyons et al. 2020; Kryazhimskiy 2021).

We also found that genetic diversity typically leads to increases in spore production of the groups. Thus, to some extent, *Dictyostelium* strains show social heterosis, at least for productivity measured as the number of spores produced (Nonacs and Kapheim 2008). This result is hard to explain unless we accept that the strains can facultatively shift their cell fate towards spore production in response to non-like cells—that is, they behave more selfishly. Other work has reached similar conclusions (Buttery et al. 2009; Madgwick et al. 2018). While a behavioral shift in response to unrelated individuals does not require active detection of foreign cells (see Parkinson et al. 2011), we know that *Dictyostelium* possesses and utilizes this ability at exactly the juncture when initial cell fates are decided (Ostrowski et al. 2008; Ho et al. 2013).

In *Dictyostelium*, kin recognition is mediated by two proteins present on the cell surface during aggregation, called TgrB1 and TgrC1 (Hirose et al. 2011). These proteins are thought to interact *in trans*, with the TgrB1 on one cell binding to the TgrC1 on a neighboring cell in a lock-and-key manner. Although the proteins were originally characterized as adhesion proteins, it is now understood that they behave as a receptor-ligand pair, with TgrC1 acting as the ligand and TgrB1 as the receptor, such that signals external to the cell are transmitted internally through TgrB1 (Hirose et al. 2017). However, little is known about how changes in TgrB1-TgrC1 binding—which would occur when strains with different *tgrB1* and *tgrC1* alleles co-aggregate—influences downstream signaling and whether these signals (or lack thereof) impact cell-fate decisions. Our results potentially indicate a reduced propensity to form stalk cells and greater selfishness when co-developing with foreign strains, but confirmation of this possibility requires further work. These findings suggest, however, that *D. discoideum*’s kin recognition genes might not only mediate avoidance of non-kin but also decisions to behave more selfishly when that avoidance is imperfect.

Our findings also differ from similar experiments in the prokaryotic organism *Myxococcus xanthus*, which also undergoes cooperative fruiting body formation in response to starvation. In that species, allopatric strains showed enhanced antagonism when co-developed to form fruiting bodies, and decreases in sporulation were also observed (Fiegna and Velicer 2005). Moreover, chimeric synergy was observed among *Myxococcus* strains that had been co-isolated from nature (Pande and Velicer 2018), suggesting local co-adaptation that is mutualistic. The differences between these two systems are striking, given the broad similarities in how they form multicellular fruiting bodies that should, in principle, be subject to the same types of conflict.

Multicellular fruiting body formation in *Dictyostelium* is a complex process that requires communication and cooperation and ultimately leads to the self-sacrifice of some of the contributing cells—and yet the overall morphology of the group is likely to confer a collective benefit, probably for spore dispersal. By exploring how group composition leads to group phenotypes, and how alterations in group phenotypes can alter the fitness benefits of collectives, we develop a better understanding of the heritability of traits at higher levels of biological organization and the potential feedbacks from individuals to groups and back again (Farine et al. 2015; Cantor et al. 2021). Addressing how and why group behaviors evolve is challenging, but the *Dictyostelium* model system offers numerous advantages. The ease of creating genetically defined groups allowed us to address many factors simultaneously, all of which potentially influence group-level phenotypes and the fitness advantages they can provide. This work points to a previously unrecognized cost of genetically diverse groups mediated via the group’s collective phenotype and demonstrates how group-level characteristics might feedback to influence selection on the individual.

## Literature Cited

Adley, K.E., Keim, M., Williams, R.S.B. 2006. Defining the genetic basis of drug action and inositol triphosphate analysis. Pp. 517–534 in E.L. Rivero F., ed. Dictyostelium discoideum Protocols. Humana Press.

Baym, M., S. Kryazhimskiy, T. D. Lieberman, H. Chung, M. M. Desai, and R. Kishony. 2015. Inexpensive multiplexed library preparation for megabase-sized genomes. PLoS One 10:e0128036.

Beerenwinkel, N., L. Pachter, and B. Sturmfels. 2007. Epistasis and shapes of fitness landscapes. Stat. Sin. 17:1317–1342.

Benabentos, R., S. Hirose, R. Sucgang, T. Curk, M. Katoh, E. A. Ostrowski, J. E. Strassmann, D. C. Queller, B. Zupan, G. Shaulsky, and A. Kuspa. 2009. Polymorphic members of the lag gene family mediate kin discrimination in Dictyostelium. Curr. Biol. 19:567–572.

Bourke, A. F. G. 2011. Principles of Social Evolution. Oxford University Press.

Broersma, C., and E. A. Ostrowski. 2022. Group transformation: fruiting body and stalk formation. P.131–150 in Herron P. L. Conlin, and W. C. Ratcliff, eds. The Evolution of Multicellularity. CRC Press.

Buttery, N. J., D. E. Rozen, J. B. Wolf, and C. R. L. Thompson. 2009. Quantification of social behavior in D. discoideum reveals complex fixed and facultative strategies. Curr. Biol. 19:1373–1377.

Cantor, M., A. A. Maldonado-Chaparro, K. B. Beck, H. B. Brandl, G. G. Carter, P. He, F. Hillemann, J. A. Klarevas-Irby, M. Ogino, D. Papageorgiou, L. Prox, and D. R. Farine. 2021. The importance of individual-to-society feedbacks in animal ecology and evolution. J. Anim. Ecol. 90:27–44.

Chen, G., G. Shaulsky, and A. Kuspa. 2004. Tissue-specific G1-phase cell-cycle arrest prior to terminal differentiation in Dictyostelium. Development 131:2619–2630.

Christensen, C., and A. N. Radford. 2018. Dear enemies or nasty neighbors? Causes and consequences of variation in the responses of group-living species to territorial intrusions. Behav. Ecol. 29:1004–1013.

Cole, B. J., and D. C. Wiernasz. 1999. The selective advantage of low relatedness. Science 285:891–893.

Crozier, R. H. 1986. Genetic clonal recognition abilities in marine invertebrates must be maintained by selection for something else. Evolution 40:1100–1101.

Farine, D. R., P.-O. Montiglio, and O. Spiegel. 2015. From individuals to groups and back: the evolutionary implications of group phenotypic composition. Trends Ecol. Evol. 30:609–621.

Fiegna, F., and G. J. Velicer. 2005. Exploitative and hierarchical antagonism in a cooperative bacterium. PLoS Biol. 3:e370.

Fortunato, A., D. C. Queller, and J. E. Strassmann. 2003. A linear dominance hierarchy among clones in chimeras of the social amoeba Dictyostelium discoideum. J. Evol. Biol. 16:438–445.

Foster, K. R., A. Fortunato, J. E. Strassmann, and D. C. Queller. 2002. The costs and benefits of being a chimera. Proc. Biol. Sci. 269:2357–2362.

Hall, A. E., K. Karkare, V. S. Cooper, C. Bank, T. F. Cooper, and F. B.-G. Moore. 2019. Environment changes epistasis to alter trade-offs along alternative evolutionary paths. Evolution 73:2094–2105.

Helanterä, H., J. E. Strassmann, J. Carrillo, and D. C. Queller. 2009. Unicolonial ants: where do they come from, what are they and where are they going? Trends Ecol. Evol. 24:341–349.

Herron, M. D., J. M. Borin, J. C. Boswell, J. Walker, I.-C. K. Chen, C. A. Knox, M. Boyd, F. Rosenzweig, and W. C. Ratcliff. 2019. De novo origins of multicellularity in response to predation. Sci. Rep. 9:2328.

Hirose, S., R. Benabentos, H.-I. Ho, A. Kuspa, and G. Shaulsky. 2011. Self-recognition in social amoebae is mediated by allelic pairs of tiger genes. Science 333:467–470.

Hirose, S., G. Chen, A. Kuspa, and G. Shaulsky. 2017. The polymorphic proteins TgrB1 and TgrC1 function as a ligand–receptor pair in Dictyostelium allorecognition. J. Cell Sci. 130:4002–4012.

Hirose, S., B. Santhanam, M. Katoh-Kurosawa, G. Shaulsky, and A. Kuspa. 2015. Allorecognition, via TgrB1 and TgrC1, mediates the transition from unicellularity to multicellularity in the social amoeba Dictyostelium discoideum. Development 142:3561–3570.

Ho, H.-I., S. Hirose, A. Kuspa, and G. Shaulsky. 2013. Kin recognition protects cooperators against cheaters. Curr. Biol. 23:1590–1595.

Hughes, W. O. H., and J. J. Boomsma. 2004. Genetic diversity and disease resistance in leaf-cutting ant societies. Evolution 58:1251–1260.

Keller, L., and M. Chapuisat. 1999. Cooperation among selfish individuals in insect societies. Bioscience 49:899–909.

Kessin, R. H. 2003. Cell motility: Making streams. Nature 422:481–482.

Kessin, R. H. 2001. Dictyostelium: Evolution, Cell Biology, and the Development of Multicellularity. Cambridge University Press.

Kryazhimskiy, S. 2021. Emergence and propagation of epistasis in metabolic networks. Elife 10.

Kuzdzal-Fick, J. J., L. Chen, and G. Balázsi. 2019. Disadvantages and benefits of evolved unicellularity versus multicellularity in budding yeast. Ecol. Evol. 9:8509–8523.

Lambrechts, L. 2010. Dissecting the genetic architecture of host-pathogen specificity. PLoS Pathog. 6:e1001019.

Lyons, D. M., Z. Zou, H. Xu, and J. Zhang. 2020. Idiosyncratic epistasis creates universals in mutational effects and evolutionary trajectories. Nat Ecol Evol 4:1685–1693.

Madgwick, P. G., B. Stewart, L. J. Belcher, C. R. L. Thompson, and J. B. Wolf. 2018. Strategic investment explains patterns of cooperation and cheating in a microbe. Proc. Natl. Acad. Sci. U. S. A. 115:E4823–E4832.

Márquez-Zacarías, P., P. L. Conlin, K. Tong, J. T. Pentz, and W. C. Ratcliff. 2021. Why have aggregative multicellular organisms stayed simple? Curr. Genet. 67:871–876.

McAuliffe, K., R. Wrangham, L. Glowacki, and A. F. Russell. 2015. When cooperation begets cooperation: the role of key individuals in galvanizing support. Philos. Trans. R. Soc. Lond. B Biol. Sci. 370:20150012.

Modlmeier, A. P., C. N. Keiser, J. V. Watters, A. Sih, and J. N. Pruitt. 2014. The keystone individual concept: an ecological and evolutionary overview. Anim. Behav. 89:53–62.

Modlmeier, A. P., J. E. Liebmann, and S. Foitzik. 2012. Diverse societies are more productive: a lesson from ants. Proc. Biol. Sci. 279:2142–2150.

Nonacs, P., and K. M. Kapheim. 2008. Social heterosis and the maintenance of genetic diversity at the genome level. J. Evol. Biol. 21:631–635.

Olson, R. S., A. Hintze, F. C. Dyer, D. B. Knoester, and C. Adami. 2013. Predator confusion is sufficient to evolve swarming behaviour. J. R. Soc. Interface 10:20130305.

Ostrowski, E. A. 2020. Evolution of multicellularity: one from many or many from one? Curr. Biol. 30:R1306–R1308.

Ostrowski, E. A., M. Katoh, G. Shaulsky, D. C. Queller, and J. E. Strassmann. 2008. Kin discrimination increases with genetic distance in a social amoeba. PLoS Biol. 6:e287.

Pande, S., and G. J. Velicer. 2018. Chimeric synergy in natural social groups of a cooperative microbe. Curr. Biol. 28:262–267.

Parkinson, K., N. J. Buttery, J. B. Wolf, and C. R. L. Thompson. 2011. A simple mechanism for complex social behavior. PLoS Biol. 9:e1001039.

Pentz, J. T., P. Márquez-Zacarías, G. O. Bozdag, A. Burnetti, P. J. Yunker, E. Libby, and W. C. Ratcliff. 2020. Ecological advantages and evolutionary limitations of aggregative multicellular development. Curr. Biol. 30:4155–4164.e6.

Poelwijk, F. J., V. Krishna, and R. Ranganathan. 2016. The context-dependence of mutations: a linkage of formalisms. PLoS Comput. Biol. 12:e1004771.

Rainey, P. B., and M. Travisano. 1998. Adaptive radiation in a heterogeneous environment. Nature 394:69–72.

Saar, M., P.-A. Eyer, T. Kilon-Kallner, A. Hefetz, and I. Scharf. 2018. Within-colony genetic diversity differentially affects foraging, nest maintenance, and aggression in two species of harvester ants. Sci. Rep. 8:13868.

Sailer, Z. R., and M. J. Harms. 2017a. Detecting high-order epistasis in nonlinear genotype-phenotype maps. Genetics 205:1079–1088.

Sailer, Z. R., and M. J. Harms. 2017b. High-order epistasis shapes evolutionary trajectories. PLoS Comput. Biol. 13:e1005541.

Sih, A., S. F. Hanser, and K. A. McHugh. 2009. Social network theory: new insights and issues for behavioral ecologists. Behav. Ecol. Sociobiol. 63:975–988.

Sih, A., and J. V. Watters. 2005. The mix matters: behavioural types and group dynamics in water striders. Behaviour 142:1417–1431.

Strassmann, J. E., and D. C. Queller. 2011. Evolution of cooperation and control of cheating in a social microbe. Proc. Natl. Acad. Sci. U. S. A. 108 Suppl 2:10855–10862.

Strassmann, J. E., Y. Zhu, and D. C. Queller. 2000. Altruism and social cheating in the social amoeba Dictyostelium discoideum. Nature 408:965–967.

Temeles, E. J. 1994. The role of neighbours in territorial systems: When are they “dear enemies”? Anim. Behav. 47:339–350.

Votaw, H. R., and E. A. Ostrowski. 2017. Stalk size and altruism investment within and among populations of the social amoeba. J. Evol. Biol. 30:2017–2030.

Wade, M. J. 2000. Epistasis as a genetic constraint within populations and an accelerant of adaptive divergence among them. Pp. 213–231 in Wolf Jason B., Brodie Iii, Edmund D. and M. J. Wade, eds. Epistasis and the Evolutionary Process. Oxford University Press.

Weinreich, D. M., Y. Lan, C. S. Wylie, and R. B. Heckendorn. 2013. Should evolutionary geneticists worry about higher-order epistasis? Curr. Opin. Genet. Dev. 23:700–707. Elsevier BV.

West-Eberhard, M. J. 1979. Sexual selection, social competition, and evolution. Proc. Am. Philos. Soc. 123:222–234.

West-Eberhard, M. J. 1983. Sexual selection, social competition, and speciation. Q. Rev. Biol. 58:155–183.

Wheatcroft, D., and T. D. Price. 2018. Collective action promoted by key individuals. Am. Nat. 192:401–414.

Wolf, J. B. 2000. Indirect genetic effects and gene interactions. Pp. 158–176 in Wolf Jason, B., Brodie Iii, Edmund, D.and M. J. Wade, eds. Epistasis and the Evolutionary Process. Oxford University Press.

