## Supplement for "From individual behaviors to collective outcomes: fruiting body formation in *Dictyostelium* as a group-level phenotype"

**Supplemental Methods:**

1. **Calculation and Interpretation of Walsh Coefficients**
2. **Illumina Genome Sequencing and Variant Calling**
3. **Calculation of Genetic Distance**
4. **Fruiting Body Imaging**

**Supplemental Results:**

**Figures S1 – S4**

**Tables S1 – S3**

### 1. Calculation and interpretation of Walsh coefficients

The application of Walsh coefficients to estimate epistatic interactions of different orders is described by Poelwijk et al (2016) and Weinreich et al. (2013, 2018). Briefly, for any set of $L$ biallelic loci (e.g., presence/absence of a particular allele, usually mutant vs wild-type), there are $\binom{L}{k}$ subsets of $k$ mutations and a total of $2^L$ possible genotypes. Each genotype is represented by a binary string of length $L$, where 0 represents the wild-type allele, and 1 represents a mutant allele at that locus. For example, if there are two alleles at each of three loci, then there are $2^3$ or 8 possible genotypes, consisting of one wild-type (000), three single mutants (001, 010, and 100), three double mutants (011, 101 and 110), and one triple mutant (111). If all possible genotypes can be assigned a fitness valued, then a Walsh-Hadamard transform can be used to generate epistasis coefficients of $k$ different orders.

Briefly, the Walsh transform, which is the digital analog of a Fourier transform, is a linear decomposition of a vector $\overrightarrow{W}$ (consisting of $2^L$ fitness estimates, representing a complete fitness landscape) into a vector of $2^L$ epistasis coefficients $\vec{E}$. The first-order coefficients ($k$=1) are proportional to classical selection coefficients and capture the fitness effect of adding a mutant allele to each possible genetic background (see Weinreich et al. 2013). Second-order coefficients account for the effect of adding a mutation dependent on the presence or absence of a mutation at a second locus, and third-order coefficients account for how pairwise interactions change dependent on the presence of a third mutation. The coefficients are orthogonal to one another (collectively, orthonormal), and therefore higher-level epistatic coefficients are independent of epistatic coefficients at lower orders. Thus, third-order coefficients capture genetic interactions *after* accounting for the effects of each mutation singly and all possible pairwise combinations. Notably, the Walsh transform can only be carried out for combinatorially complete datasets—that is, estimation of Walsh coefficients requires fitness information on all possible combinations of a set of alleles.

To date, the Walsh transform has been applied to examine pairwise and higher-order epistasis among different mutations when added to a focal genetic background. For example, Poelwijk et al (2019) used the Walsh transform to explore pairwise and higher-order epistasis among 13 mutations (resulting in analysis of $2^{13}$ or 8,192 genotypes) in a fluorescent protein of *Entacmaea quadricolor*. Lozovsky et al (2021) compared the adaptive landscape of resistance to five different antimalarial drugs by constructing the 16 possible combinations of four amino acid substitutions in the dihydrofolate reductase gene of *Plasmodium falciparum*. The adaptive landscapes for four of the drugs showed sub-optimal fitness peaks (i.e., ruggedness generated by epistasis), but the topologies of the landscapes were similar among the different drugs and higher-level epistasis (>2$^{nd}$ order) did not contribute strongly to the adaptive landscape.

The above studies have used the method of estimating Walsh coefficients to assess epistasis among mutations within a genome and its impact on the phenotype of a population of cells. However, epistasis can arise at multiple levels of biological organization: within genomes, between genomes (e.g., nuclear mitochondrial interactions) and between individuals of the same or different species, referred to as 'genotype-by-genotype' epistasis or G ×G (Wolf 2000; Linksvayer 2007; Buttery et al. 2010; Heath 2010; Teseo et al. 2014; Walsh et al. 2020; Turkarslan et al. 2021). In the case of social interactions, genotype-by-genotype epistasis for the group phenotype can arise when the phenotypic effect on an allele in one individual depends on the alleles present in social partners. The presence of social epistasis has important consequences for the evolutionary trajectories of social traits, as the genetic context-dependence can potentially lead to co-adaptation and coevolution, and runaway processes, such as arms races and potentially speciation (Linksvayer 2007).

Although a number of studies have examined epistasis among genotypes with respect to social behavior (Linksvayer 2007; Buttery et al. 2010; Saltz 2013; Walsh et al. 2020), there has so far been no application of Walsh coefficients to examine the contribution of higher-order (>2$^{nd}$ order) epistasis to group outcomes. The *Dictyostelium* model system, however, is particularly amenable to this type of analysis because it is feasible to create many groups of defined genetic composition and to measure the outputs of those groups repeatedly and over short periods of time. In addition, because the phenotype of interest here (sporulation) is intrinsically a higher-level phenotype, it is amenable to this type of analysis. In other words, we are not measuring the sporulation of each individual in response to genotypes of social partners (i.e., indirect genetic effects) but exploring the group phenotype as a function of the genetic composition of the group.

**2. Illumina Genome Sequencing and Variant Calling**

*Isolation of Genomic DNA* - We grew cells of EO189, EO192, EO607, EO610, EO621, EO622, and EO623 in tissue culture plates by inoculating 7 mL of HL5 with glucose (Formedium) containing 2X PSV (10 µg/ml penicillin, 50 µg/ml streptomycin sulfate, 60 ng/ml vitamin $B_{12}$, and 20 ng/ml folate), 1% fetal bovine serum, and 1% autoclaved *K. pneumoniae* with spores, changing this medium to HL5 containing 2X PSV (see below) and 1% FBS the next day, and washing the cells off the plate with KK2 on the third day. We heat-shocked spores of EO195, EO347, EO350, and EO354 to obtain cells (30 min at 45ºC, 5 min at 4ºC) and incubated the solution overnight on a platform shaker in KK2 buffer, which was supplemented with 2X PSV for the latter three strains. We used a 1.4X or higher ratio of Ampure XP beads to sample volume ratio to purify the EO195 and EO608 gDNA samples and a 1.5X ratio of Zymo Select-a-size DNA clean and concentrator MagBeads for EO621, EO622, EO623, EO347, EO350, and EO354.

To obtain high molecular weight genomic DNA, we followed a modified version of Adley and colleagues' (2006) protocol. Specifically, we added 0.05 mg RNase A to the initial DNAzol solution and incubated the resuspended pellet for 35 min. We precipitated the DNA with an equal volume of 100% ethanol and pelleted the DNA for 10 minutes. We resuspend the DNA pellet in 30 µl of 8 mM NaOH and adjusted the pH to 8.0 by adding 3 µl of 0.1 M HEPES. We extracted EO608 genomic DNA from spores that were incubated in DNAzol and RNase A for an hour.

*Library preparation and Illumina sequencing* - For Illumina library preparation, we used the Nextera XT Library Prep Kit Reference Guide (Illumina Document # 15031942 v03), with quarter-sized reactions for EO189, EO192, EO607, and EO610. We modified the protocol by using 0.26 ng/ul gDNA for EO195 and EO608 and twice the recommended concentration of normalized gDNA (0.4 ng/µl) for the remaining strains. For EO347, EO350, and EO354, we improved amplification by reducing the elongation temperature in the Nextera XT Amplify Libraries thermal-cycler program to 65°C, increasing this part of the cycle to 60 seconds (Su et al. 1996), and increasing the number of cycles from 12 to 14. For EO621, EO622, and EO623, we used KAPA HiFi HotStart ReadyMix (2X) instead of Nextera PCR Master Mix, with 25 µl reactions and ran a thermal-cycler program as described in Baym et al. (2015).

We followed the Nextera XT Clean Up Libraries protocol and used a 0.67X AMPure XP beads to library volume ratio for VA and NC samples. For the MA and VA samples, we used a 0.64X SPRI Select beads ratio for the MA and VA samples and reduced the pellet air drying time to 5-7 minutes. We eluted the libraries in 15 µl RSB (resuspension buffer) and recovered 13-14 µl of each library. To measure each library concentration, we used a ThermoFisher Qubit 2.0 flourometer, and we had the UH Seq-N_Edit Core (SNEC) determine fragment sizes with an Agilent 4200 TapeStation or Agilent 2100 BioAnalyzer. We sent pools of "libraries with a concentration of 4 nM (10-20 uL)" or a more dilute equivalent to Admera Health in South Plainfield, NJ for sequencing on an Illumina HiSeq X Ten Platform.

*Fastq alignment and variant calling* - Illumina reads were mapped to the genome of AX4, the standard *D. discoideum* reference genome (downloaded from dictybase.org; GFF3 files dated November 30, 2016). Prior to mapping, we masked a chromosome 2 duplication specific to this lab strain (chromosome DDB0232429 from positions 3016082 to 3768654). To map reads to the reference, we used bwa-mem (0.7.17-r1188) -M. We used picard 2.20.2-0 to sort the resulting sam files and convert them to bam files with SortSam SORT_ORDER=coordinate, MarkDuplicates, and then BuildBamIndex. After running gatk (gatk4-4.0.12.0-0) HaplotypeCaller --emit-ref-confidence GVCF, we used gatk (gatk4-4.1.3.0-0) CombineGVCFs to combine the g.vcfs of all 12 strains. We ran gatk (gatk4-4.1.3.0-0) GenotypeGVCFs -all-sites -new-qual on the final combined samples to obtain a multisample vcf ready for filtering.

*SNP Calling* - We used bcftools (bcftools 1.9) norm -d none --threads 6 to remove duplicate entries from our multi-sample vcf file. We then used a hard filter in gatk (gatk4-4.1.3.0-0) VariantFiltration, using a modified haploid version of their generic recommendations: --filter-expression "QD < 2.0", "FS > 60.0", "MQ < 40.0", "SOR > 3.0." To remove non-variable sites and further filter out alternative variants with low coverage, we ran gatk VariantFiltration --genotype-filter-expression "isHomVar == 1 && DP < 4.0". For some down-stream analyses it was necessary to remove variants that did not pass the filters from the vcf, so we used gatk SelectVariants --exclude-filtered to do so. We subset our vcf to include the six *D. discoideum* chromosomes using bcftools view --regions. We also ran gatk SelectVariants --restrict-alleles-to BIALLELIC and removed sites with missing data with vcftools --max-missing 1.

**3. Calculation of Genetic Distances**

We used vcfR (1.10.0) to read the multisample vcf into R (3.6.2) and convert it into a genlight object with the vcfR2genlight function. We used the poppr (2.8.6) as.snpclone wrapper to convert this into a snpclone object for calculating Euclidean distance with the poppr bitwise.dist function (euclidean = TRUE). To create the unrooted tree in Fig. 1 C, we used the ape (5.3) plot function (typ="unrooted", show.tip=TRUE) to plot the neighbor joining tree produced by the ape nj function that we ran with Euclidean distances.

### 4. Fruiting Body Imaging

We imaged the fruiting bodies from above and from the side for three blocks of our NC/MA and VA/NH experiments. Briefly, we imaged the filters at 7.81X magnification on a Leica M205FA stereomicroscope with a color temperature of 2900K from the KL 1500 LCD light source. The overhead images were taken with a monochrome camera with an 87 ms exposure, whereas the images from the side were taken in color with a 137 ms exposure. The following macro was run in Image J (2.0.0-rc-69/1.52k) to analyze the overhead images:

```
run("Set Scale...", "distance=142 known=1 pixel=1 unit=mm");
makeRectangle(819,402,912,933);
run("Crop");
run("8-bit");
run("Threshold...");
setAutoThreshold("Default dark");
setOption("BlackBackground", false);
run("Make Binary", "thresholded remaining black");
run("Watershed");
run("Analyze Particles...", "size=0.005-0.2 circularity=0.75-1.00 show=Outlines display exclude
summarize");
```

To analyze the size of the fruiting bodies from the side images of filters, we first manually cropped the images at the base of the stalk in Image J. We then used the following macro in batch mode to estimate sorus height for all images. This process locates the sorus (ball of spores) and then measures the distance from the sorus to the base of the fruiting body.

```
run(“Set Scale…”, “distance=142 known=1 pixel=1 unit=mm”);
makeRectangle(852.3, 0, 852.3, 822);
run(“Crop”);
run(“8-bit”);
run(“Invert”);
run(“Threshold Regional Gradient”, “circularity=0.6 minimum=80 maximum=3600
fill_phase_detected display method=Slow”);
run(“Make Binary”);
```

```
run(“Watershed”);
rename(x[2]);
run(“Set Measurements…”, “centroid invert display add redirect=None decimal=3”);
run(“Analyze Particles…”, “size=0.01-0.25 circularity=0.74-1.00 show=Outlines display exclude summarize record”);
Table.rename(“Results”, x[2]);
saveAs(“Results”, dir + name + “.csv”);
```

**Supplemental Figures**

**Figure S1.** Example overhead images of fruiting bodies for a set of NC/MA mixes. Numbers indicate the mix (e.g., ‘12’ is the mix of strains 1 and 2).

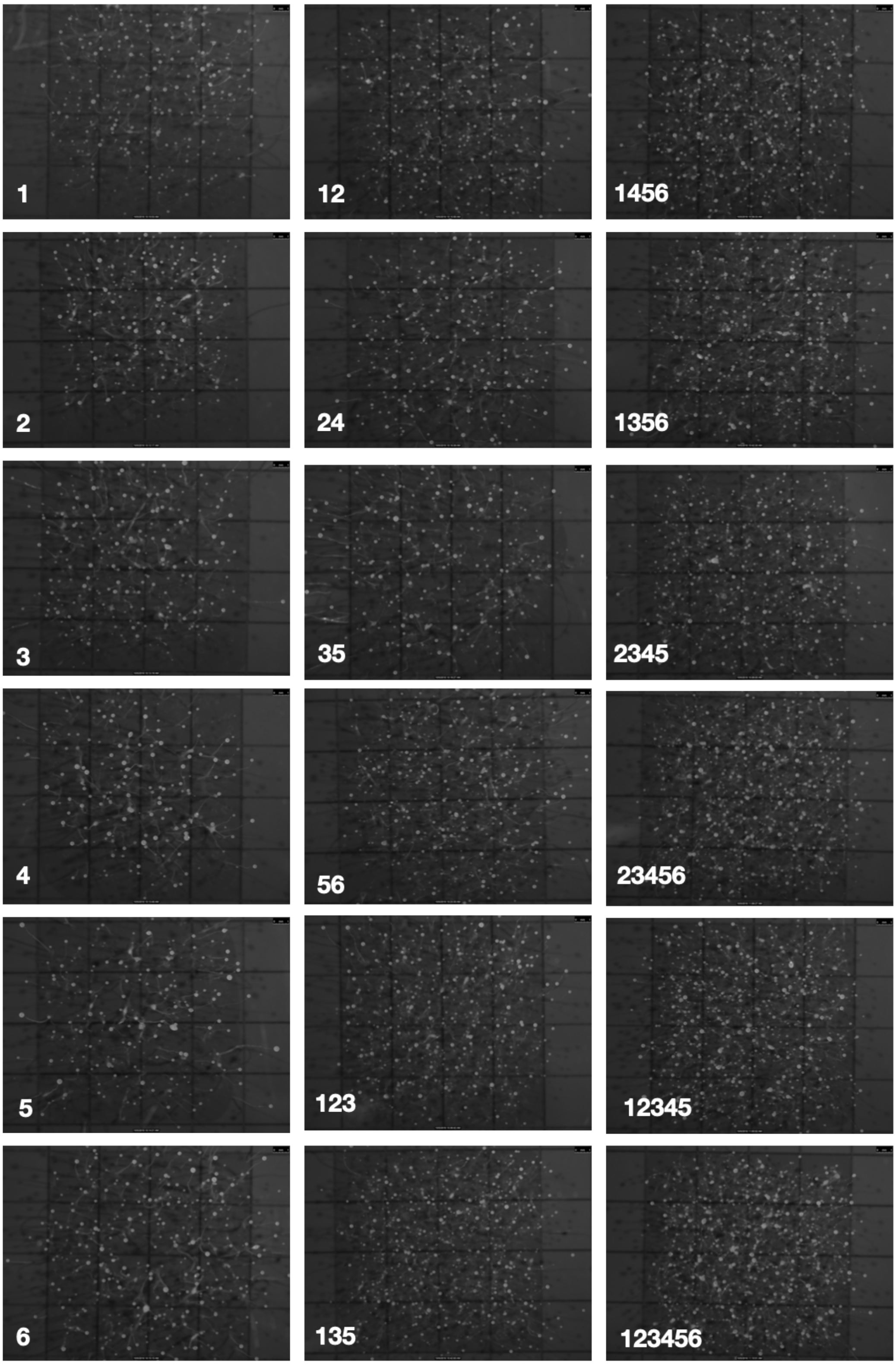

**Figure S2.** Example side images of fruiting bodies for a set of NC/MA mixes. Numbers indicate the mix (e.g., '12' is the mix of strains 1 and 2.).

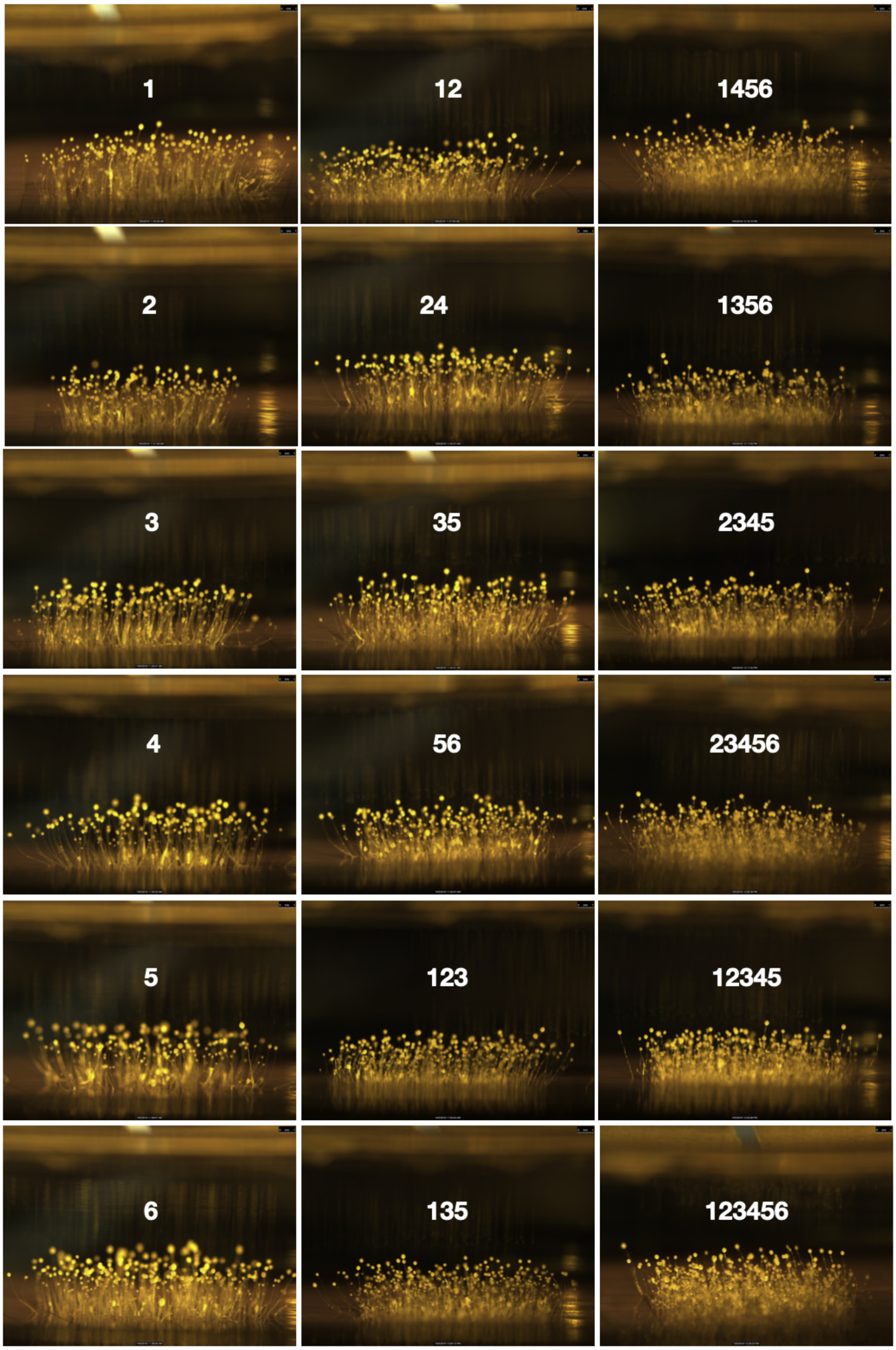

**Figure S3. Focal strain impacts on sporulation of their groups. (A)** NC/MA mixes. **(B)** VA/NH mixes. The data in each subplot (Strain 1 - Strain 6) are the same (= deviation from expected vs expected sporulation), but the points are colored differently. In each plot, blue points show mixes with a given focal strain, whereas gray points indicate mixes without the focal strain. ANCOVA was used to ask whether observed sporulation of groups that contain the focal strain was significantly greater or less than groups that lack that strain. Only strain 4 (NH) shows evidence of being a 'bad apple'— its presence is consistently associated with the group performing worse than expected in terms of sporulation. Strains 1 and 2 (from NC), strain 1 (from VA), and strain 6 (from MA), show evidence of improving the sporulation of the group. See Table S4 for ANCOVA results.

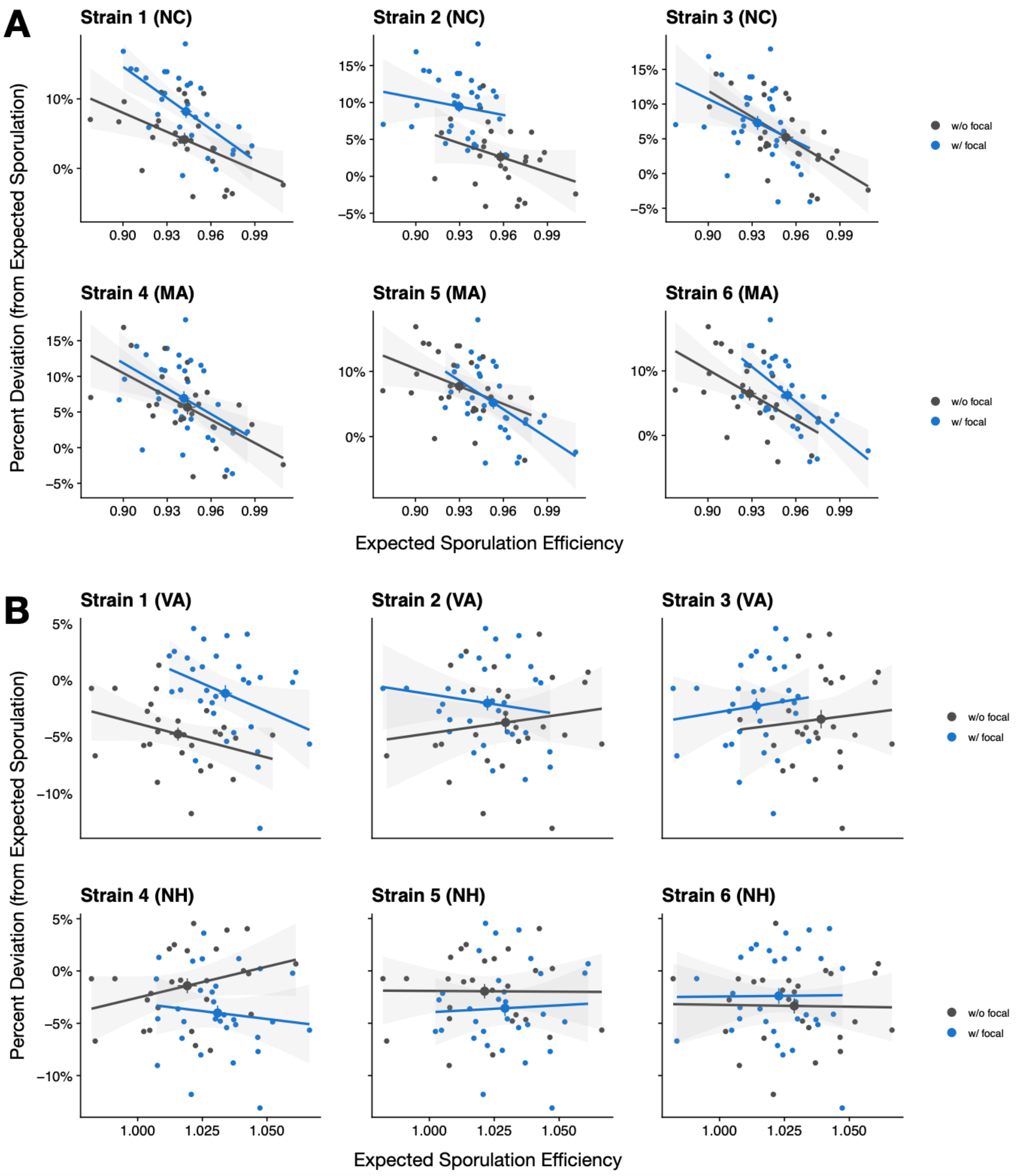

**Figure S4. Gene expression changes in response to being surrounded by nonself cells.** Hirose et al. (2015) determined which genes show altered expression in response to being surrounded by nonself (i.e., cells with different *tgrB1*-t*grC1* alleles). We merged these results with the classification of genes according to their degree of prespore or prestalk bias (Parikh et al. 2010). Genes that are downregulated in response to nonself at 12 hr (when initial cell fate is adopted) are strongly prestalk-biased (purple).

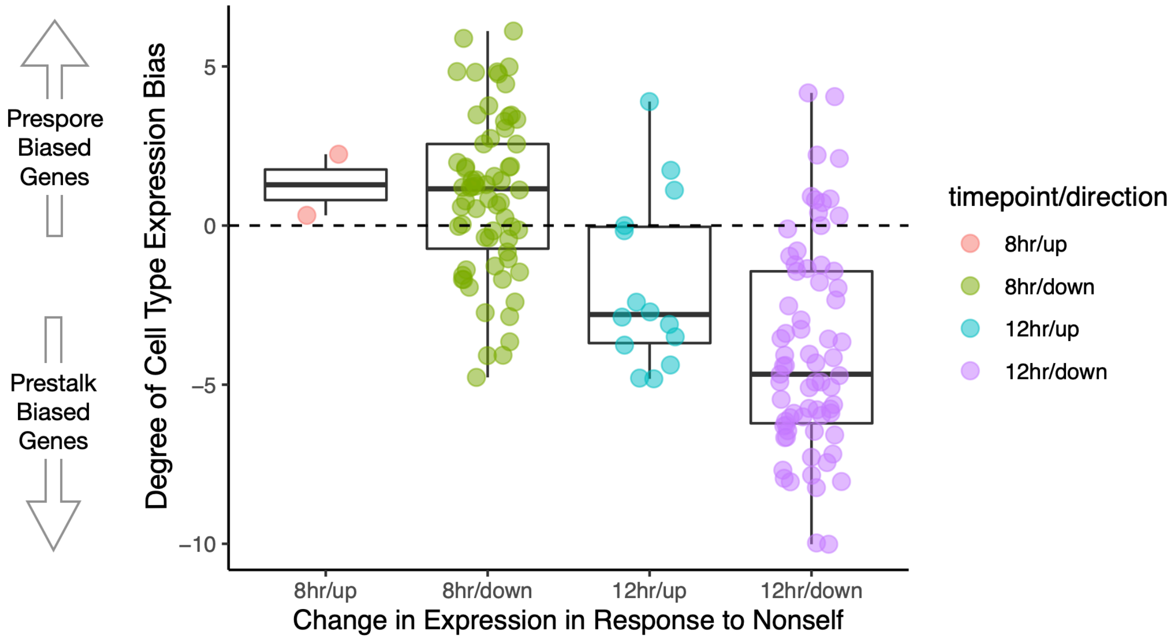

### Supplemental Tables

**Table S1. Strains used in the study and their locations of origin.** Strain ID indicates the numeric shorthand for a particular strain. For example, the VA/NH mix called "123" is the three-way mix of strains EO189, EO192, and EO195. Origin is abbreviated as follows: MLBS = Mountain Lake Biological Station, VA; NHPA = New Hampshire, Proctor Academy; GSNP = Great Smoky National Park, NC; SMFS = Smith MacLeish Field Station, MA.

| Strain | PopPair (Strain ID) | Origin | GPS Coordinates |
|---|---|---|---|
| EO189 | VA/NH (1) | MLBS | 37.35358, -80.53615 |
| EO192 | VA/NH (2) | MLBS | 37.35358, -80.53615 |
| EO195 | VA/NH (3) | MLBS | 37.35358, -80.53615 |
| EO347 | VA/NH (4) | NHPA | 43.45207, -71.825785 |
| EO350 | VA/NH (5) | NHPA | 43.45207, -71.825785 |
| EO354 | VA/NH (6) | NHPA | 43.45207, -71.825785 |
| EO607 | NC/MA (1) | GSNP | 35.60852, -83.44736 |
| EO608 | NC/MA (2) | GSNP | 35.60852, -83.44736 |
| EO610 | NC/MA (3) | GSNP | 35.60852, -83.44736 |
| EO621 | NC/MA (4) | SMFS | 42.448581, -72.681296 |
| EO622 | NC/MA (5) | SMFS | 42.448581, -72.681296 |
| EO623 | NC/MA (6) | SMFS | 42.448581, -72.681296 |

**Table S2. Bonferroni-adjusted confidence intervals for the deviation from expected sporulation by sites (sympatric mixes only).** 95% confidence interval for the intercept for NC strains (top row) and the three other sites, considering sympatric mixes only (i.e., two- or three-way mixes where all strains come from the indicated site). Observed sporulation of groups was modeled as a function the expected sporulation, number of strains (2 or 3), and site. *P*-values, adjusted for multiple comparisons, indicate that NC strains are significantly different from all the rest (all $P$<0.0001; adj method=Holm) and thus that their sporulation as a group is significantly more positive than expected. All other sites (NH, VA, and MA) are not significantly different from each other.

| | 2.50% | 97.50% |
|---|---|---|
| (Intercept) | 0.106 | 0.506 |
| Expected spores | 0.609 | 0.933 |
| Number of strains | -0.017 | 0.069 |
| site VA | -0.191 | -0.079 |
| site NH | -0.236 | -0.124 |
| site MA | -0.191 | -0.058 |

| | NC | VA | NH | |
|---|---|---|---|---|
| NC | - | | | |
| VA | <0.0001 | - | | |
| NH | <0.0001 | 0.13 | - | |
| MA | <0.0001 | 0.65 | 0.45 | - |

**Table S3. Linear model assessing how genetic diversity, genetic or geographic relationships influence how well strains co-sporulate.** Significance tests based on Type III sums of squares using Anova() in the car package.

| | df | SS | *F* | *P* (>F) |
|---|---|---|---|---|
| Expected Sporulation | 1 | 3.88 | 511.6 | <0.0001 |
| Population pair (NC/MA or VA/NH) | 1 | 0.81 | 107.4 | <0.0001 |
| Number of Strains | 1 | 0.33 | 43.0 | <0.0001 |
| Genetic distance (Euclidean) | 1 | 0.20 | 26.9 | <0.0001 |
| Allopatric/Sympatric | 1 | 0.07 | 9.3 | 0.002 |
| Block | 1 | 0.06 | 7.3 | 0.007 |
| Population pair × block | 1 | 0.20 | 26.6 | <0.0001 |
| Residuals | 505 | 3.83 | | |

**Table S4. 'Bad apple' analyses:** *P*-values associated with main effect of strain on the group's deviation from expected spores. ANCOVA with expected sporulation as a predictor as well as strain-by-expected interaction. Asterisks indicate ANCOVA significance: * $P<0.05$, ** $P<0.01$, *** $P<0.001$.

| Strain | Strain-by-expected interaction | Strain effect (ANCOVA) | |
|---|---|---|---|
| Strain 1 (NC) | 0.202 | <0.001 | *** |
| Strain 2 (NC) | 0.616 | <0.001 | *** |
| Strain 3 (NC) | 0.673 | 0.9 | |
| Strain 4 (MA) | 0.844 | 0.439 | |
| Strain 5 (MA) | 0.331 | 0.914 | |
| Strain 6 (MA) | 0.359 | 0.009 | ** |
| Strain 1 (VA) | 0.514 | <0.001 | *** |
| Strain 2 (VA) | 0.268 | 0.112 | |
| Strain 3 (VA) | 0.923 | 0.181 | |
| Strain 4 (NH) | 0.142 | 0.01 | * |
| Strain 5 (NH) | 0.814 | 0.131 | |
| Strain 6 (NH) | 0.922 | 0.384 | |

### References

Adley, K.E., Keim, M., Williams, R.S.B. 2006. Defining the genetic basis of drug action and inositol triphosphate analysis. Pp. 517–534 *in* E. L. Rivero F., ed. *Dictyostelium discoideum* Protocols. Humana Press.

Baym, M., S. Kryazhimskiy, T. D. Lieberman, H. Chung, M. M. Desai, and R. Kishony. 2015. Inexpensive multiplexed library preparation for megabase-sized genomes. PLoS One 10:e0128036.

Buttery, N. J., C. R. L. Thompson, and J. B. Wolf. 2010. Complex genotype interactions influence social fitness during the developmental phase of the social amoeba *Dictyostelium discoideum*. J. Evol. Biol. 23:1664–1671.

Heath, K. D. 2010. Intergenomic epistasis and coevolutionary constraint in plants and rhizobia. Evolution 64:1446–1458.

Hirose, S., B. Santhanam, M. Katoh-Kurosawa, G. Shaulsky, and A. Kuspa. 2015. Allorecognition, via TgrB1 and TgrC1, mediates the transition from unicellularity to multicellularity in the social amoeba *Dictyostelium discoideum*. Development 142:3561–3570.

Linksvayer, T. A. 2007. Ant species differences determined by epistasis between brood and worker genomes. PLoS One 2. Public Library of Science.

Lozovsky, E. R., R. F. Daniels, G. D. Heffernan, D. P. Jacobus, and D. L. Hartl. 2021. Relevance of Higher-Order Epistasis in Drug Resistance. Mol. Biol. Evol. 38:142–151.

Parikh, A., E. R. Miranda, M. Katoh-Kurasawa, D. Fuller, G. Rot, L. Zagar, T. Curk, R. Sucgang, R. Chen, B. Zupan, W. F. Loomis, A. Kuspa, and G. Shaulsky. 2010. Conserved developmental transcriptomes in evolutionarily divergent species. Genome Biol. 11:R35.

Poelwijk, F. J., V. Krishna, and R. Ranganathan. 2016. The context-dependence of mutations: a linkage of formalisms. PLoS Comput. Biol. 12:e1004771.

Poelwijk, F. J., M. Socolich, and R. Ranganathan. 2019. Learning the pattern of epistasis linking genotype and phenotype in a protein. Nat. Commun. 10:4213.

Saltz, J. B. 2013. Genetic composition of social groups influences male aggressive behaviour and fitness in natural genotypes of Drosophila melanogaster. Proceedings of the Royal Society B: Biological Sciences 280:20131926. Royal Society.

Su, X. Z., Y. Wu, C. D. Sifri, and T. E. Wellems. 1996. Reduced extension temperatures required for PCR amplification of extremely A+T-rich DNA. Nucleic Acids Res. 24:1574–1575.

Teseo, S., N. Châline, P. Jaisson, and D. J. C. Kronauer. 2014. Epistasis between adults and larvae underlies caste fate and fitness in a clonal ant. Nat. Commun. 5:3363.

Turkarslan, S., N. Stopnisek, A. W. Thompson, C. E. Arens, J. J. Valenzuela, J. Wilson, K. A. Hunt, J. Hardwicke, A. L. G. de Lomana, S. Lim, Y. M. Seah, Y. Fu, L. Wu, J. Zhou, K. L. Hillesland, D. A. Stahl, and N. S. Baliga. 2021. Synergistic epistasis enhances the co-operativity of mutualistic interspecies interactions. ISME J., doi: 10.1038/s41396-021-00919-9.

Walsh, J. T., A. Garonski, C. Jackan, and T. A. Linksvayer. 2020. The collective behavior of ant groups depends on group genotypic composition.

Weinreich, D. M., Y. Lan, J. Jaffe, and R. B. Heckendorn. 2018. The influence of higher-order epistasis on biological fitness landscape topography. J. Stat. Phys. 172:208–225.

Weinreich, D. M., Y. Lan, C. S. Wylie, and R. B. Heckendorn. 2013. Should evolutionary geneticists worry about higher-order epistasis? Curr. Opin. Genet. Dev. 23:700–707.

Wolf, J. B. 2000. Gene interactions from maternal effects. Evolution 54:1882–1898.
